# Predicting binding events in very flexible, allosteric, multi-domain proteins

**DOI:** 10.1101/2024.06.02.597018

**Authors:** Andrea Basciu, Mohd Athar, Han Kurt, Christine Neville, Giuliano Malloci, Fabrizio C. Muredda, Andrea Bosin, Paolo Ruggerone, Alexandre M. J. J. Bonvin, Attilio V. Vargiu

## Abstract

Knowledge of the structures formed by proteins and small molecules is key to understand the molecular principles of chemotherapy and for designing new and more effective drugs. During the early stage of a drug discovery program, it is customary to predict ligand-protein complexes *in silico*, particularly when screening large compound databases. While virtual screening based on molecular docking is widely used for this purpose, it generally fails in mimicking binding events associated with large conformational changes in the protein, particularly when the latter involve multiple domains. In this work, we describe a new methodology to generate bound-like conformations of very flexible and allosteric proteins bearing multiple binding sites by exploiting only information on the unbound structure and the putative binding sites. The protocol is validated on the paradigm enzyme adenylate kinase, for which we generated a significant fraction of bound-like structures. A fraction of these conformations, employed in ensemble-docking calculations, allowed to find native-like poses of substrates and inhibitors (binding to the active form of the enzyme), as well as catalytically incompetent analogs (binding the inactive form). Our protocol provides a general framework for the generation of bound-like conformations of challenging drug targets that are suitable to host different ligands, demonstrating high sensitivity to the fine chemical details that regulate protein’s activity. We foresee applications in virtual screening, in the prediction of the impact of amino acid mutations on structure and dynamics, and in protein engineering.

## Introduction

Molecular recognition is a fundamental process for cellular life, regulation, and pathology,^1^ yet its quantitative understanding remains a major challenge due to the complexity of accounting for interactions among flexible partners fluttering in a crowded solution. The structural determinants of molecular recognition are best described indeed considering an ensemble of conformational states of each (macro)molecule involved.^1,2^ Among them, proteins represent the majority of interactors and span a very wide flexibility spectrum, ranging from side-chain reorientations to large-scale domain motions, possibly coupled to secondary structure variations.^1^ Target plasticity is key especially for multi-specific proteins, whereby even minor conformational changes can enable the binding of multiple compounds to different regions of the same broad binding site.^3–5^

The rapid increase in the number of experimentally resolved protein structures within the last decades has fueled the development of computational tools to mine their conformational space, including machine/deep-learning approaches integrating experimental data and simulations.^6–25^ Methods to mimic protein-ligand association *in silico*, such as molecular docking and virtual screening, have become routinary in any modern drug design lab.^1,26,27^ Indeed, predicting the interactions between proteins and small molecules (ligands) underlies modern chemotherapy and drug design.^1,28,29^ Despite its widespread use, the proper description of partners’ flexibility in molecular docking remains a big challenge in the field, significantly affecting accuracy.^1,26,28,30^ This difficulty is primarily due to the challenge of exploring plasticity in high-dimensional spaces, coupled with well-known sensitivity of docking to even minor structural changes at the binding interface.^1^ These difficulties affect also AI-based algorithms, particularly for the generation of protein ensembles of high conformational diversity or in the presence of interactions with membranes,^31^ although recent algorithms achieved state-of-the-art accuracy in reproducing protein-ligand structures, at least for common natural compounds.^32,33^

Among the strategies developed to cope with the flexibility issue,^28^ ensemble-docking accounts for plasticity by using a pre-defined set of structures of one (generally the protein) or both the binding partners.^26,34,35^ The method has been successfully applied to various targets, showcasing its versatility in drug discovery efforts.^29,36–38^ Within this framework, we recently proposed EDES (Ensemble Docking with Enhanced sampling of pocket Shape),^39–41^ a method employing metadynamics simulations^42–46^ with a set of *ad hoc* collective variables to bias the shape and the volume of a binding site. EDES was validated on a set of non-allosteric globular proteins bearing a single binding site, enabling the prediction of their bound(holo)-like conformations. Here we largely re-designed the original protocol, hereafter referred to as generalized EDES (gEDES), to deal with multi-domain allosteric proteins bearing extended binding sites composed of multiple (sub)pockets. These proteins include indisputably relevant and challenging targets for drug design efforts.^47,48^

We validated gEDES on the pharmaceutically important enzyme adenylate kinase (ADK), a paradigm protein undergoing very large structural rearrangements upon binding of multiple substrates to an extended region composed of two (sub)pockets (**Figure 1**).^49,50^ Structurally, ADK is as a monomeric enzyme composed of a main domain (CORE) linked to a NMP-binding (NMP) and an ATP-binding (LID) domain.^49^ These three domains embed two distinct binding regions at the interfaces between the CORE and the NMP and the LID regions (hereafter NC and LC respectively in **Figure 1**), which bind ATP and AMP (substrates for the phosphoryl transfer) or two ADP molecules during the reverse reaction. During the catalytic cycle, the NMP and LID domains close over the substrate(s) via hinge-like motions (pink arrows in **Figure 1**),^51–53^ generating a reactive environment shielded from non-structural water molecules.^54,55^

**Figure 1.**
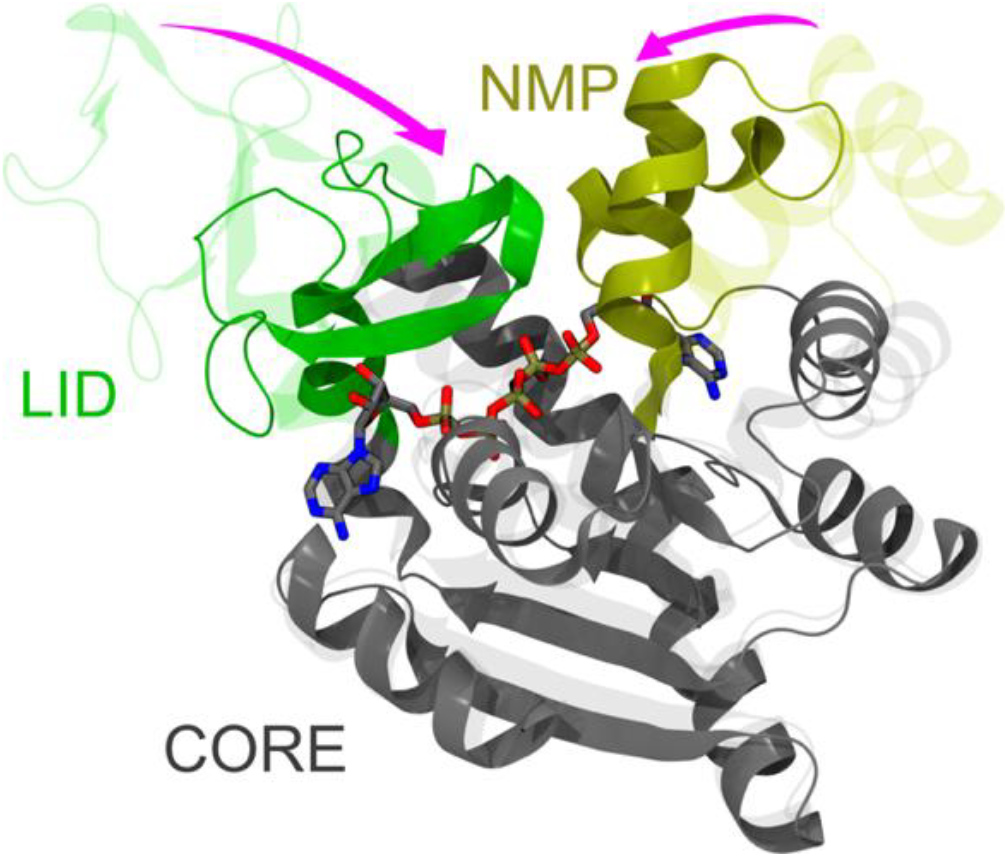
Comparison between the apo (PDB ID: 4AKE) and holo (PDB ID: 1AKE) experimental structures of ADK, represented by transparent and solid ribbons, respectively. The protein structure is built upon three quasi-rigid domains, called CORE, LID and NMP, and it features two distinct binding sites. The LID (residues 118–160) and NMP (residues 30–61), which undergo hinge-like motions upon binding of substrates or inhibitors, are colored green and dark yellow, respectively. The CORE domain (residues 1–29, 62–117, 161–214) is colored gray. The inhibitor P^1^,P^5^-bis(adenosine-5^’^-)pentaphosphate (AP5) is shown by sticks colored by atom type (C, O, N, P atoms in grey, red, blue, and tan, respectively). The main hinge-like conformational changes between adjacent quasi-rigid domains are highlighted by the pink arrows.

This behavior is shared by different classes of pharmaceutically relevant proteins, including kinases and transferases.^56,57^ Allosteric models of ADK activity have been proposed on the basis of the high correlation between the structural rearrangements of the LC and NC interfaces.^58–60^ While cofactors such as Mg^2+^ are essential for the catalysis,^61,62^ substrate binding and related conformational changes are believed to be largely independent of their presence.^61,63^

Due to its peculiarities, ADK has been the subject of several experimental^63–67^ and computational^58,68–83^ studies shedding light on the details of the structural rearrangements and the energetics governing its biological activity. Moreover, ADK has been extensively employed as paradigm system to benchmark various computational strategies aiming to reproduce large/allosteric apo(open)/holo(closed) conformational transitions,^70,71,75–78,84–89^ as well as in molecular docking studies.^80,90^

Here, we demonstrate that our new protocol can generate a significant fraction of structures of ADK that are very similar to those bound to a set of different ligands, without exploiting any information about these compounds. Importantly: i) the ligands include substrates and inhibitors bound to a closed (active) conformation of the enzyme, as well as an incompetent binder bound to an open (inactive) state; ii) the agreement encompasses the fine geometry of the extended binding region, resulting in the correct side-chain orientation of most residues therein. Moreover, when employed in ensemble-docking calculations, these conformations: i) yielded native-like poses of substrates and inhibitors of ADK among the top-ranked ones for all the ligands; ii) reproduced the binding mode of the catalytically inactive GTP analog to an open structure of ADK. Our findings highlight the high accuracy of gEDES in accounting for the fine chemical details regulating enzyme activity (a feature that AI-based methods could not easily catch, as demonstrated below) and place it among the state-of-the-art tools for accurate *in silico* Structure-Based Drug Design.

## Results and discussion

The workflow of gEDES is sketched in **Figure 2**. The only two ingredients needed to the method are the unbound structure of a protein and the location of its putative binding sites (BSs). Next, a dissection of the protein into quasi-rigid (QR) domains linked by flexible hinges is performed, and the collective variables (CVs) to perform bias-exchange well-tempered metadynamics^42–44^ simulations are setup. These are the gyration radius of the BS (RoG_BS_), the three Contacts across Inertia Planes (CIP_1-3_), and the “contacts between quasi-Rigid Domains” (cRD) defined between residues belonging to adjacent QR domains and lining the BS (here the LID-CORE and NMP-CORE interfaces – hereafter LC and NC, respectively; see Methods for implementation details). If, as in the case of ADK, a significant fraction of residues defining the cRDs are charged, the corresponding CV is further split into one containing only the charged residues and another one containing all remaining ones. As demonstrated below, such a splitting allows for effective sampling of sidechain conformations of the former group, which otherwise could be penalized by the lower free energy barriers associated with large fluctuations of non-charged residues. Once the simulations are done, a multi-step cluster analysis is performed on the resulting trajectories, producing several structure representatives to be employed in ensemble docking calculations.

**Figure 2.**
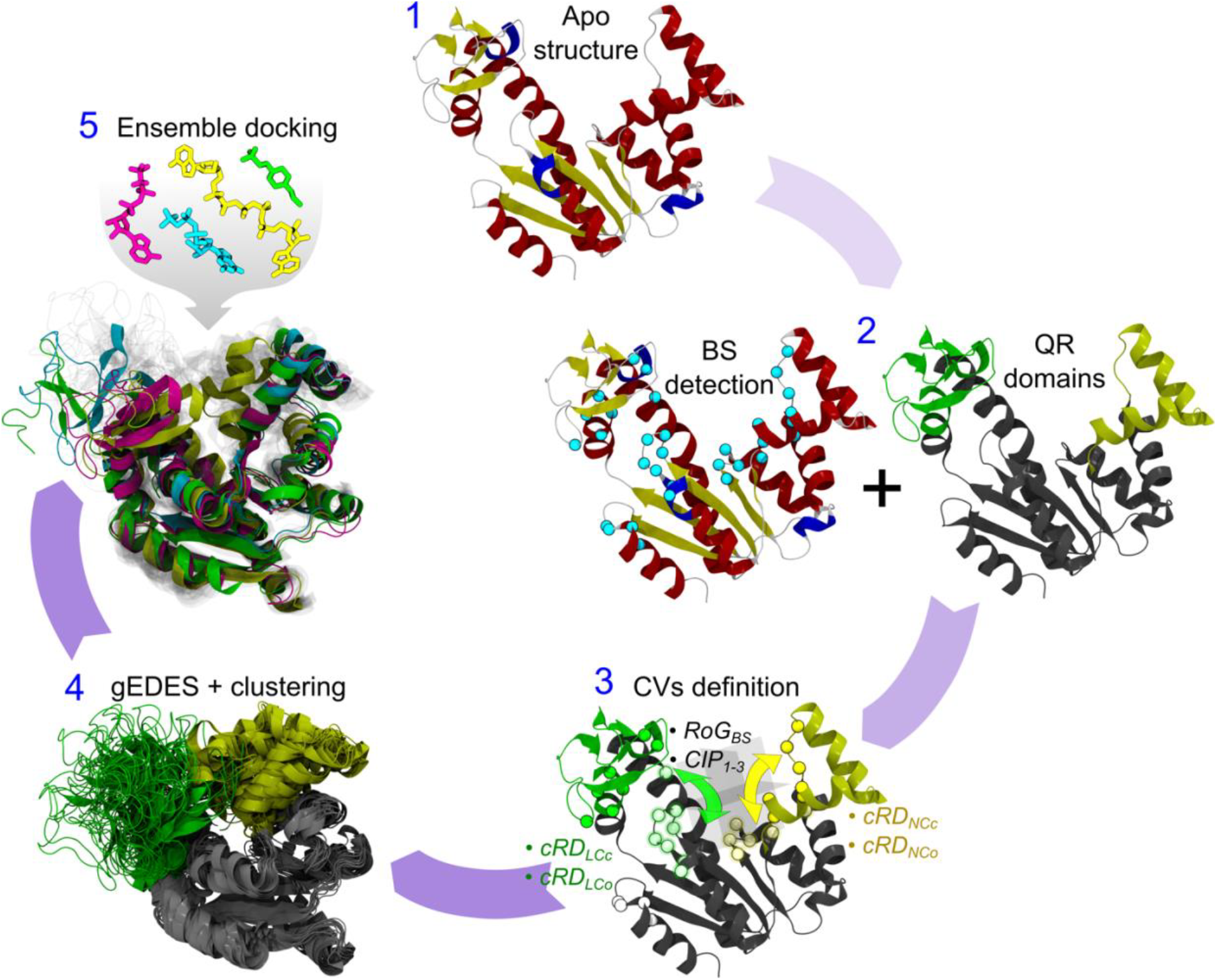
gEDES workflow. After choosing a structural template of the apo structure of the protein (1) either from experiments or from modelling, a list of putative binding site(s) and quasi-rigid (QR) domains linked by flexible hinges is identified by exploiting experimental information or computational algorithms such as COACH-D. (2). Next, the collective variables (CVs) are setup (3): in addition to the gyration radius (RoG_BS_) and the three contacts across inertia planes (CIP_1-3_), a set of “contacts between quasi-Rigid Domains” (cRD) is also defined between residues belonging to adjacent QR domains and to the BS (here the LID-CORE and NMP-CORE interfaces – hereafter LC and NC, respectively; see Methods for implementation details). If, as for ADK, a relevant fraction of residues lining the cRDs features a charged sidechain, this CV is further split into two new ones containing, respectively, only the charged residues and all the remaining ones. Finally, a multi-step cluster analysis is performed (4) on the trajectory generated with the gEDES setup, producing several structure representatives to be employed in ensemble docking calculations (5).

In the following, we report the performance of gEDES in reproducing the structures of ADK bound to the four ligands employed in this work, focusing on the fine geometry of the BSs. Next, we show how accounting for protein plasticity improves docking outcomes. The experimental closed (active) structures of ADK in complex with substrates ADP and AMP (PDB IDs: 1ANK,^54^ 2ECK^91^) and the inhibitor P^1^,P^5^-bis(adenosine-5^’^-)pentaphosphate (hereafter AP5; PDB ID: 1AKE^92^), as well as the open (inactive) structure in complex with the non-hydrolysable (catalytically incompetent) GTP analog β,γ-methyleneguanosine 5′-triphosphate (herafter GCP; PDB ID: 6F7U^63^), are used as a reference for validation. Note that the structures of ADK bound to ADP and AMP ligands feature an overall conformation very similar to the one bound to AP5, with sub-angstrom structural variations at the corresponding BSs (see **Table S5**). Therefore, in the following, our discussion will often refer only to the AP5 and GCP bound structures.

### Sampling of holo-like protein conformations

The performance of gEDES in generating holo-like structures of ADK is summarized in **Figure 3** and **Table 1**. Notably, the BS employed to define the CVs (that is the one found by COACH-D, hereafter referred to as BS_COACH_; see Methods for details) does not coincide precisely with that of any ligand (hereafter BS_AP5_, BS_ADP_, BS_AMP_, and BS_GCP_). Indeed, BS_COACH_ is much larger than all of them but BS_AP5_, and as such it is not biased towards specific compounds; it comprises 32 residues, 11 of which (almost 35% of the total) are charged. By inspecting the RMSD distributions in the left column of **Figure 3** we see that, for all BS investigated in this work, gEDES generates a non-negligible fraction of holo-like structures. The same performance was not obtained during multiple independent standard MD simulations (hereafter MD_std_) for which, as expected, we found a consistent percentage of conformations with RMSD values lower than 2.5 Å only when using BS_GCP_ as reference. This is not surprising, as GCP is bound to an open inactive structure of the enzyme, while the binding of all the other compounds is associated with a decrease of the RoG of the corresponding BS by more than 20% (**Table S4**).

**Figure 3.**
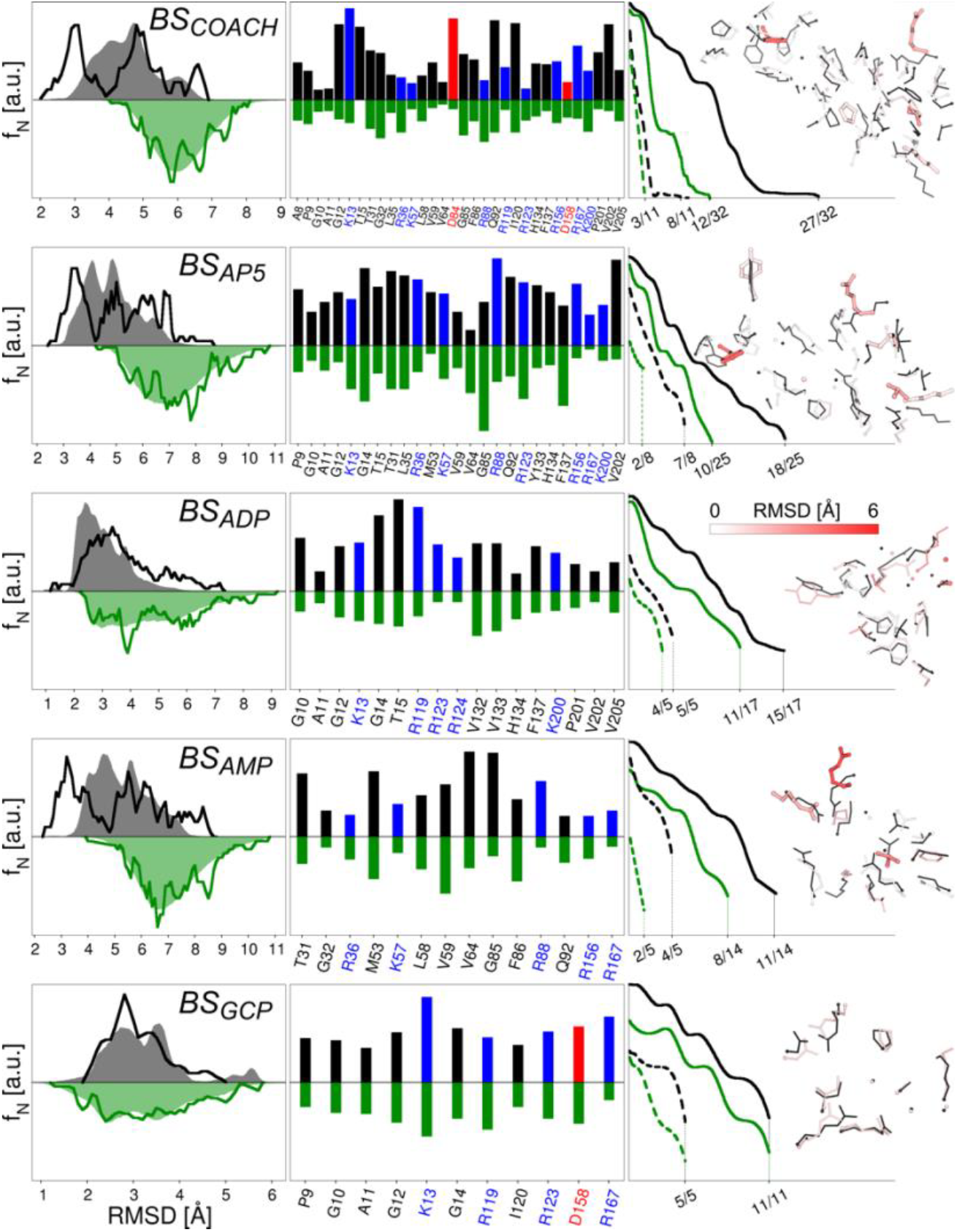
Performance of MDstd and gEDES simulations in reproducing holo-like conformations of the computationally derived (BSCOACH, first row) and of the ADK ligands’ binding sites (2nd to last rows). The left column reports the normalized frequency distributions of the RMSD values of each site calculated on the non-hydrogenous atoms with respect to the reference structure (1AKE for BSCOACH and BSAP5, 2ECK for BSADP, 1ANK for BSAMP, 6F7U for BSGCP) after alignment of the same site. Green and gray shaded areas (lines) refer to the distributions extracted from the production trajectory (cluster representatives used for docking calculations) of the unbiased and gEDES simulations, respectively. These distributions were obtained by grouping RMSD values into bins of 0.2 Å in width and interpolating the resulting distribution with cubic splines and a density of 20 points per Å. The middle column reports the frequency of holo-like conformations sampled by each residue lining the corresponding BS during the unbiased and gEDES simulations. Bars pointing upwards refer to gEDES and are colored according to the amino acid type (red: negatively charged; blue: positively charged; black: neutral), while green bars pointing downwards refer to the unbiased simulation. A holo-like conformation is counted when the RMSD of the residue (calculated on all non-hydrogenous atoms after alignment of the whole BS) is lower than the arbitrary threshold defined for each amino acid in **Table S6** (and always lower than 2.5 Å). The right column reports the normalized distributions of the fraction of total (solid lines) and charged (dashed lines) residues simultaneously assuming a holo-like conformation. The best conformation from the ensemble of clusters is also reported against the corresponding reference structure. Sidechains are shown by sticks colored from white to red according to the value of the per-residue RMSD from the reference (thin black sticks), with thick sticks identifying those residues that do not sample holo-like conformations.

**Table 1.** Percentage of structures with RMSD values below 2 and 2.5 Å, calculated respectively on the C_α_ atoms for the protein and on the non-hydrogenous atoms for each BS. For each region values were obtained after the alignment to the same region on the reference structure. Results refer to the cumulative trajectories and to cluster representatives of both the standard MD simulation (MD_std_) and the gEDES approach. The PDB codes of each reference holo-structure are reported in the first row. The lowest values of the RMSD (Å) are reported in parentheses for each entry.

|  | Holo: 1AKE |  |  | Holo: 2ECK | Holo: 1ANK | Holo: 6F7U |  |
| --- | --- | --- | --- | --- | --- | --- | --- |
| RMSD [Å] | BS<sub>COACH</sub> | BS<sub>AP5</sub> | Protein (C<sub>α</sub>) | BS<sub>ADP</sub> | BS<sub>AMP</sub> | BS<sub>GCP</sub> | Protein (C<sub>α</sub>) |
| MD<sub>std</sub> | 0 (3.7) | 0 (3.9) | 0 (2.9) | 0.9 (2.1) | 0 (3.1) | 23.5 (1.1) | 16.2 (0.9) |
| gEDES | 1.3 (2.0) | 0.7 (2.0) | 0.3 (1.8) | 13.4 (0.9) | 2.3 (2.2) | 19.8 (1.3) | 10.1 (1.0) |
| MD<sub>std</sub><sup>clust</sup> | 0 (3.9) | 0 (4.0) | 0 (3.1) | 1.8 (2.1) | 0 (3.8) | 24.2 (1.1) | 16.8 (1.5) |
| gEDES<sup>clust</sup> | 4.0 (2.1) | 3.8 (2.2) | 0.8 (2.1) | 13.6 (1.1) | 1.5 (2.2) | 16.7 (2.0) | 3.8 (2.0) |

**Table 1** quantifies these results in terms of RMSD values from the true holo complexes for all the BSs and for the whole protein (in the case of AP5 and GCP), considering both the entire trajectory and cluster-derived structures. We observe that, while gEDES was able to sample conformations displaying respectively RMSD of as low as 2.0 (1.8) Å for BS_COACH_ (whole protein), the best structures sampled by MD_std_ featured corresponding values of 3.7 and 2.9 Å respectively. These findings clearly indicate that, even in absence of any bias on the whole protein, our set of local CVs was able to drag the entire ADK structure toward holo-like geometries. More importantly, our multi-step clustering pipeline allowed us to preserve these geometries within a restricted pool of selected structures. Indeed, the sets of conformations selected from gEDES MDs for the ensemble docking of AP5, ADP, AMP, and GCP include geometries 2.2, 1.1, 2.2, 2.0 Å away from the corresponding complex structures. As expected, gEDES and MD_std_ have similar sampling performances only in the case of BS_GCP_. These results further confirm the general applicability of gEDES, despite the protocol has been originally developed to address targets undergoing large conformational changes.

The second column in **Figure 3** reports the percentage of conformations displaying *N* amino acids *simultaneously* in their bound-like conformation (with *N* going from 0 to the number of residues lining each BS; see also **Table S6**). gEDES was able to generate a geometry of BS_COACH_ simultaneously displaying 27 out of 32 residues (of which 8 out of 11 charged amino acids) with a bound-like conformation. In turn, this resulted in the accurate reproduction of bound-like conformations for all experimental BSs, which are partly overlapping and/or included in BS_COACH_. In contrast, the best conformations generated from MD_std_ displayed less than 40% of the residues in a bound-like geometry for all binding sites but BS_GCP_ (for which the two protocols perform similarly). These different performances of gEDES and MD_std_ trace back to their diverging abilities in reproducing the bound-like conformations of *individual residues* lining BS_COACH_ and all the experimental BSs (**Figure S1**).

### Reproducing ligand native poses

We performed ensemble-docking calculations on the conformational clusters extracted from gEDES and MD_std_ trajectories using HADDOCK^93^ and AutoDock4.^94^ The poses reported for AutoDock4 were obtained after performing a ligand cluster analysis on the top poses obtained from each individual docking run (i.e. a run for each protein conformation), while in HADDOCK all cluster conformations within each ensemble were used in a single docking run. Importantly, in the spirit of using our protocol for prediction purposes, docking of all ligands was performed on the region encompassing BS_COACH_, despite this site is much larger than all the experimental sites but BS_AP5_. Details of this implementation can be found in the *Methods* section.

**Table 2** shows that for all ligands but ADP, only the conformations of ADK obtained with gEDES enabled to retrieve top-ranked native-like complex structures (the top native-like complex structures obtained from the combined set of docking calculations using HADDOCK and AutoDock4 are shown in **Figure 4**).

**Table 2.** Performance of AutoDock4 and HADDOCK in reproducing the experimental structures of the complexes between ADK and the four ligands investigated in this work. For each compound, two cluster sets (gEDES and MD_std_) were employed in ensemble-docking calculations. The Autodock results refer to clusters of docking poses obtained from a cluster analysis performed on all generated complexes (see Methods), while for the HADDOCK they refer to the single structures from a single run starting from the ensemble of MD conformations. The sampling performance is calculated as the percentage of poses within 2.5 Å from the native structure (for HADDOCK calculated at the rigid-body stage out of 50000 poses). The pose ranking refers to the first native-like conformation with the highest score within each cluster. The next row reports the heavy atoms RMSD of the ligand and the binding site, calculated for the first native-like conformation according to ranking. F_nat_ indicates the fraction of native contacts recovered within a shell of 5 Å from the ligand in the experimental structures. The overall best poses are highlighted in bold and shown in **Figure 4**.

|  |  | AP5 |  | AMP |  | ADP |  | GCP |  |
| --- | --- | --- | --- | --- | --- | --- | --- | --- | --- |
|  |  | MD<sub>std</sub> | gEDES | MD<sub>std</sub> | gEDES | MD<sub>std</sub> | gEDES | MD<sub>std</sub> | gEDES |
| Autodock | Sampl. perf. [%] | 0.6 | 1.8 | - | 0.9 | 1.9 | 2.7 | 1.2 | 1.8 |
|  | Pose rank | 2 | 2 | - (63) | **7** | 10 | 1 | 16 | **10** |
|  | RMSD<sub>lig/BS</sub> [Å] | 2.3/6.1 | 2.2/2.2 | -<br>(3.7/4.9) | **2.1/3.1** | 2.3/3.8 | 0.9/1.1 | 2.5/1.7 | **1.8/2.3** |
|  | F<sub>nat</sub> | 0.58 | 0.79 | - | 0.74 | 0.57 | 0.86 | 0.75 | 0.91 |
| HADDOCK | Sampl. perf. [%] | 2.8 | 0.57 | - | 0.004 | 0.06 | 0.1 | 1.01 | 1.4 |
|  | Pose rank | 61 | **1** | 643 | 193 | 23 | **1** | 390 | 11 |
|  | RMSD<sub>lig/BS</sub> [Å] | 2.2/7.9 | **1.6/2.1** | 2.4/7.8 | 1.7/2.7 | 2.3/4.2 | **0.6/1.4** | 2.0/3.5 | 1.7/2.0 |
|  | Best RMSD<sub>lig</sub> [Å] / rank | 1.8/68 | 0.9/16 | 2.4/643 | 1.7/193 | 2.2/81 | 0.5/7 | 1.9/644 | 0.8/74 |
|  | F<sub>nat</sub> | 0.38 | 0.82 | 0.33 | 0.67 | 0.54 | 0.88 | 0.43 | 0.86 |

**Figure 4.**
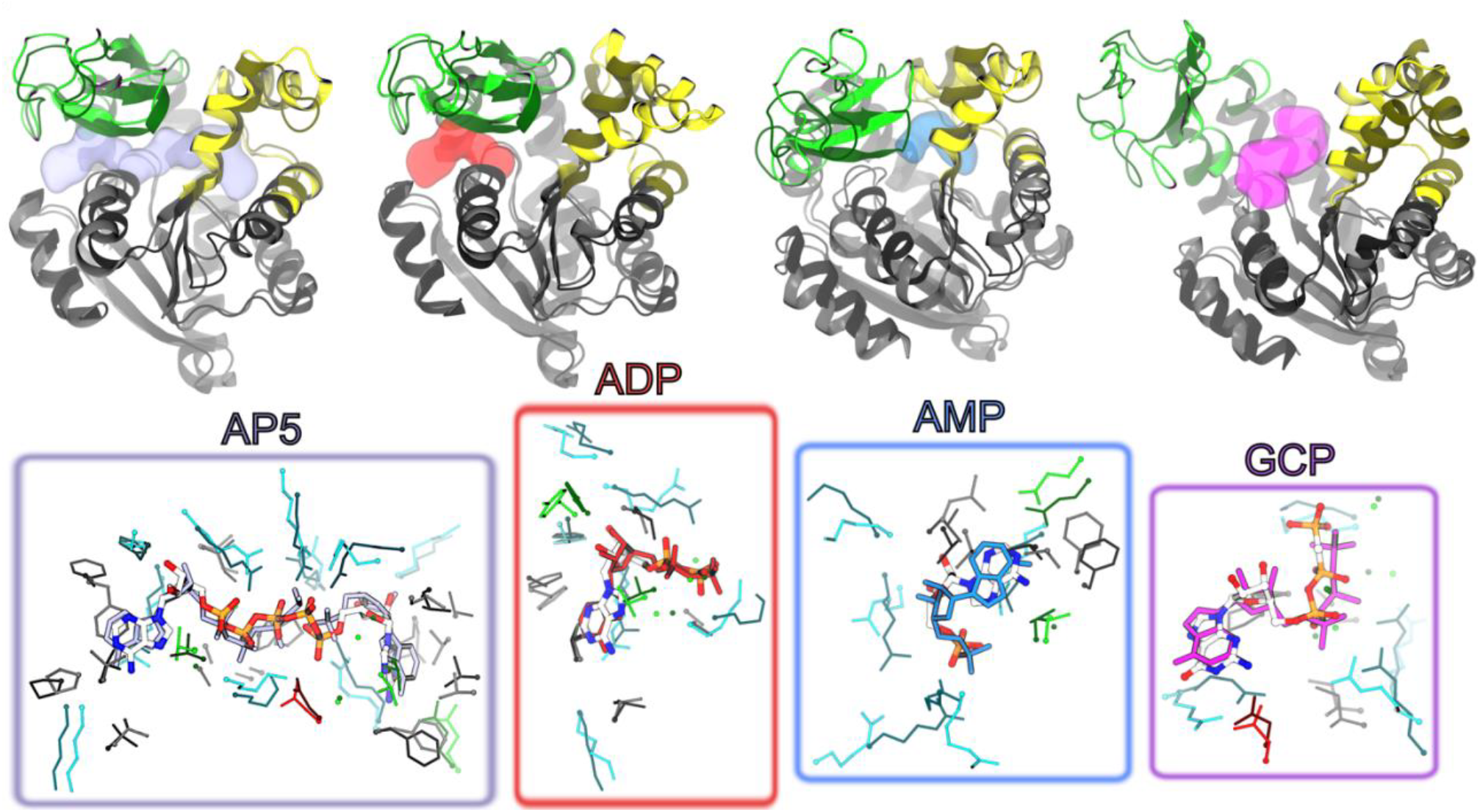
Best complex structures (nearest-to-native) obtained from the combined set of docking calculations using HADDOCK and AutoDock4 for all ADK ligands investigated in this work. These structures correspond to the best overall poses reported in **Table 2**. For each ligand (names are indicated in the figure), the top row shows the structure obtained by gEDES yielding the nearest-to-native complex geometry superposed to the corresponding experimental bound conformation. These structures are shown in cartoons and colored respectively by domain (green, yellow, and gray for the LID, NMP, and CORE domains respectively) and in dark gray. The location of the ligand in the experimental structures is shown by a semi-transparent surface colored in iceblue, red, cyan, and magenta, respectively. The bottom row shows the comparison between the geometries of the BSs and of the ligand in the experimental bound structure vs. the nearest-to-native pose. The sidechains of residues lining the BS in the former (latter) structure are shown by dark thin (light thick) sticks colored by residue type; the ligand is shown with single color sticks in the experimental structure, while thicker sticks colored by atom type (C, N, O, P atoms in white, blue, red, and orange respectively) indicate the best docking pose.

Using either AutoDock4 or HADDOCK with the clusters obtained from gEDES, the top two poses of AP5 include a near-native complex structure (RMSD_lig_ values of 2.2 and 1.6 Å, respectively); importantly, top ligand poses were found within conformations of BS_AP5_ that resembled the experimental complex, enabling to recover a large fraction of native contacts (F_nat_). On the other hand, clusters obtained from MD_std_, even if able to yield near-native ligand poses, were ranked poorly and moreover were obtained on largely distorted BS structures (RMSD_BS_ values of 4.9 Å and 7.9 Å respectively), thus describing incorrect binding modes with low F_nat_ values.

A similar situation is seen for ADP, for which both docking programs find a native-like pose (associated with low RMSD_lig_ and high F_nat_ values) ranked as first only when using gEDES clusters; native-like poses found using the MD_std_ clusters were again ranked poorly and associated with distorted BS conformations. Regarding GCP, Autodock and HADDOCK found the native-like pose ranked as 1^st^ and 11^th^ respectively, when using gEDES clusters, compared to ranks of 16^th^ and 390^th^ when using structures from MD_std_. In this case, as expected, near-native ligand poses were found on holo-like BS_GCP_ conformations with both the MD_std_ and the gEDES clusters. Finally, both docking programs retrieved near-native ligand poses of AMP on top when using gEDES cluster representatives, although these structures featured a slightly distorted binding region compared to the previous ligands and were ranked poorly by HADDOCK. On the other hand, no native-like binding mode was retrieved when using protein structures derived from MD_std_. The slightly worst gEDES performance for AMP is perhaps expected, as this compound binds to the NC interface, whose competent conformation is triggered via an allosteric boost involving a previous closure of the LID domain.^58^ Nonetheless, binding-prone conformation of BS_AMP_ enabling to accurately predict near-native AMP poses were recovered even without explicitly mimicking allosteric regulation.

### Comparison with previous works

In this section, we compare our results with (several) previous works aiming to explore ADK functional motions, eventually leading to the generation of druggable structures, and to characterize the energetics of its apo-to-holo transitions.

Flores and Gerstein employed ADK as test system for their “conformation explorer” algorithm.^95^ Based on the identification of protein’s hinge axes followed by Euler rotations and MD simulations, the authors addressed the LID motion, for which the closest-to-holo generated model displayed a C_α_-RMSD (after superimposing the CORE and NMP domains) of 3.8 Å, to be compared with the value of ∼17 Å (after the same superposition) between the experimental apo and holo structures. Krüger *et al*. applied the NMsim^70^ conformational search algorithm, based on elastic network models (ENM), to generate ADK bound-like structures either by unbiased simulations or by biasing the RoG of the protein to values below RoG_apo_. These approaches produced, respectively, conformations featuring backbone RMSD as low as 3.06 Å and 2.36 Å from the holo structure.^51,92^ A similar methodology was employed by Ahmed *et al*.,^71^ who coupled a rigid cluster normal-mode analysis (RCNMA) with NMsim to generate conformations featuring a C_α_-RMSD as low as 1 or 3.1 Å from the experimental complex depending on whether the ADK closed or open structures were used. Unfortunately, no information was reported by the authors on the performance in reproducing bound-like geometry of the BS.

Wang *et al*.^96^ employed Replica-Exchange MD simulations (starting from the holo structure) to estimate the free energy profile and the timescales associated with LID opening and closing. They found these values to be respectively ∼29 μs and ∼118 μs, in good correlation with the experimental data^88^ and pointing to the need for very long, unbiased simulations to collect a statistically significant number of opening/closing events. Yasuda *et al*.^75^ employed “parallel cascade selection MD (PaCS-MD)” to generate transition pathways between the (known) apo and holo (AP5-bound complex) structures of ADK. The method produced models displaying a C_α_-RMSD as low as 1.1 Å from the target structure, although also in this case no data regarding the BS geometry were reported. Jalalypour and coworkers^76^ developed an approach to identify key residues responsible for specific conformational transitions in proteins by comparing apo and holo structures of a generic target. Steered MD (SMD)^58^ simulations fed with this information were employed to trigger apo-to-holo functional rearrangements in ADK, generating conformations with C_α_-RMSD of 3.1 Å from the AP5-bound geometry.

A thorough comparison of our method with those discussed above is limited by the lack of data regarding the accuracy in reproducing the geometry of the extended binding region considered in the present work, which is relevant for drug design applications. Below, we report three examples enabling (at least in part) such an assessment.

In the first one, Kurkcuoglu and Doruker^80^ included ADK in the set of 5 proteins selected to assess the performance of their ENM-based workflow to generate an ensemble of holo-like protein conformations for docking calculations. Their best model (selected after filtering the RoG of the protein so as to discard conformations with values larger than RoG_apo_) displayed a C_α_-RMSD of 2.4 Å over the whole protein.^51,92^ When these models were employed in docking calculations of AP5, the closest-to-native pose featured an RMSD_lig_ 2.9 Å from the experimental structure. In the second example, we employed NeuralPLexer,^32^ a recent computational approach exploiting deep learning to predict protein-ligand complex structures by integrating small molecules information and biophysical inductive bias, to reproduce the structures of the complexes between ADK and the four ligands investigated in this work. We used as inputs the unbound structure of the protein and the conformations of the ligands extracted from the corresponding complexes. Finally, we also employed AlphaFold3^33^ to predict the structures of the complexes formed by ADK with ADP and AMP. In this case, the user provides the sequence of the protein and selects the ligand from a list of available compounds.

The results of these calculations, summarized in **Table 3**, indicate that gEDES (a purely biophysical method) compares well with NeuralPLexer and AlphaFold3. While the latter displays the best performance for ADP and AMP (common natural compounds that are highly represented in the PDB database), it was not possible to evaluate its performance for AP5 and GCP, which are not included in the dataset of ligands. Interestingly, NeuralPLexer predicts complex structures that are slightly closer than those generated by gEDES and Autodock/HADDOCK for AP5 and AMP, while it has a slight worst performance for ADP. Moreover, it fails in generating native-like conformations for GCP.

**Table 3.** Comparison of gEDES, NeuralPLexer, and AlphaFold3 performances in reproducing native-like conformations of the complexes between AK and the four ligands investigated in this work. See caption of **Table 2** for further details.

|  |  | AP5 | AMP | ADP | GCP |
| --- | --- | --- | --- | --- | --- |
| Autodock-gEDES | RMSD<sub>lig/BS</sub> [Å] | 2.2/2.2 (2) | 2.1/3.1 (7) | 0.9/1.1 (1) | 1.8/2.3 (10) |
|  | F<sub>nat</sub> | 0.79 | 0.74 | 0.86 | 0.91 |
| HADDOCK-gEDES | RMSD<sub>lig/BS</sub> [Å] | 1.6/2.1 (1) | 1.7/2.7 (193) | 0.6/1.4 (1) | 1.7/2.0 (11) |
|  | F<sub>nat</sub> | 0.82 | 0.67 | 0.88 | 0.86 |
| NeuralPLexer | RMSD<sub>lig/BS</sub> [Å] | 1.6/1.9 (1) | 1.3/2.1 (1) | 1.0/1.8 (1) | -* |
|  | F<sub>nat</sub> | 0.89 | 0.90 | 0.77 | -* |
| AlphaFold3 | RMSD<sub>lig/BS</sub> [Å] | -<sup>†</sup> | 0.9/1.3 (1) | 0.3/0.6 (1) | -<sup>†</sup> |
|  | F<sub>nat</sub> | -<sup>†</sup> | 1 | 1 | -<sup>†</sup> |

\*NeuralPLexer was unable to correctly reproduce the conformation of the protein and of the BS (minimum RMSD ~5.9 Å), as well as native-like poses of GCP (RMSD<sub>min</sub> ~ 3.8 Å, F<sub>nat</sub> = 0.50)
<sup>†</sup>The AlphaFold3 server (https://alphafoldserver.com/) allows in the current implementation to predict the binding of only a few ligands, including AMP and ADP.

It is instructive to comment on the results for AMP and GCP in view of the published data on the biological function of ADK. A recent investigation^67^ suggested that initial binding of AMP followed by ATP could lead to a closed state that does not allow for the correct positioning of the two ligands for effective phosphate transfer. In other words, binding of AMP should occur after binding of the other substrate for optimal enzymatic activity; therefore, our results are remarkable as we did not perform any prior simulation of the AMP-ATP complex. Even more interesting are the results obtained for GCP, whose binding arrests the enzyme in a catalytically non-functional open state. Since our approach generates widely different conformations, including some open structures, we were able to retrieve native-like conformations of the complex between GCP and ADK. In contrast, NeuralPLexer generated in all cases catalytically competent structures very similar to those found for the true substrates of the enzyme. We hypothesize that this could be due, at least partly, to the large predominance of substrate bound (closed) ADK conformations. In the future, it would be instructive to assess the performance of NeuralPLexer, AlphaFold3 and similar methodologies on datasets including active and inactive compounds binding to the same protein in different conformations.

## Concluding remarks and perspectives

The accurate prediction of the molecular determinants enabling biological and therapeutic activity mediated by proteins is a holy grail in computational biology and drug design. Accuracy and computational costs are two essential factors to be accounted for in developing effective software and protocols. The first factor requires not only to identify the correct structures of protein-ligand complexes, but also to distinguish between active and inactive compounds based on their binding modes and affinities. This is particularly relevant when dealing with enzymes, whose selectivity is critical for the precise control of metabolic pathways often occurring in crowded environments filled with many chemically related substrates.

In this work, we proposed and validated gEDES, a computational protocol for the accurate prediction of holo-like conformations of proteins, including allosteric and/or multi-pocket targets undergoing extended conformational changes upon ligand binding. Notably, gEDES relies only on the knowledge of a structure of the protein (even if unbound to any ligand) and on the identification of its putative binding site(s), thereby avoiding biases towards any specific chemotype. We validated our methodology on the paradigm enzyme adenylate kinase, widely employed to benchmark computational methods aiming to reproduce apo/holo conformational transitions and to predict holo-like conformations of proteins. gEDES was able to generate a large fraction of holo-like conformations of both the whole protein and the extended binding competent region, which is constituted by multiple distinct binding sites. Importantly, the agreement goes beyond the identification of the overall correct geometry of the binding region, defined by its backbone atoms, and includes reproduction of holo-like sidechains conformations. Furthermore, these binding competent geometries were reproduced for both active and inactive protein conformations within a single simulation. When using a limited set of conformations extracted from gEDES trajectories in ensemble-docking calculations of substrates, inhibitors, and catalytically incompetent binders of adenylate kinase, we retrieved in all cases native-like structures of the complexes (which include closed and open protein conformations) ranked among the top models. These results demonstrate that gEDES, coupled to state-of-the-art docking simulations, can achieve very high sensitivity towards subtle but crucial chemical details of ligands, placing it among the state-of-the-art methodologies in the field.

In perspective, we plan to use our protocol to predict functional conformational changes in proteins, in accurate virtual screening campaigns, and in the rational design and repurposing of drugs. Exploring the conformational diversity of binding sites in a “chemotype-unbiased” manner could lead to the identification of a larger number of promising lead candidates^1,97–100^ and/or to discover potential new uses for already marketed ones.^101^ In addition, the detection of putative binding regions could in principle be improved by combining existing structural data with druggability estimations based on the physico-chemical properties of the site.^21,102^ Sampling could be further improved by introducing new CVs crucial for protein dynamics and/or coupling metadynamics with other enhanced-sampling methods. For example, biasing the radius of gyration of the whole protein could be of help in case of large proteins, for which exploiting only variables defined on a putative binding region could not be effective. Another possibility is the coupling of our strategy with others directly addressing the fine treatment of torsional angles^103^ rotations and/or secondary structure changes.^104^ Our workflow also allows for the inclusion of experimental information at different stages of the process, for instance during post-processing as done e.g. in ref. 80. For instance, the cluster analysis can be biased to extract protein conformations featuring a value for the radius of gyration within a desired range. Finally, accurate structural data generated by gEDES simulations can be exploited to improve the predictivity of AI-based methods. In this respect, a strong limitation of these methods is that they predict only static structures and not dynamical ensembles, which should closely reproduce the behavior of biomolecular systems. As a long-term purpose, we aim to create a database of protein structures encompassing bound, unbound, and intermediate conformational states.

## Methods

### Binding site determination

In this work, we employed a bacterial ADK enzyme to validate the protocol. The binding region of ADK was identified by uploading the apo structure with PDB ID: 4AKE^105^ (resolved at 2.2 Å resolution and devoid of any missing residue) to the webserver COACH-D.^106^ COACH-D gives in output a set of up to 10 putative binding sites, together with a ranking C-score ranging from 0 (unreliable prediction) to 1 (highly accurate prediction). In the case of ADK, the software identified three binding regions (**Table S1**) displaying a C-score respectively of 0.99 (BS1_COACH_), 0.89 (BS2_COACH_), and 0.67 (BS3_COACH_), while the remaining predictions featured very low C-scores (<0.01) and as such were discarded. For our purposes, we identified a consensus binding region (hereafter BS_COACH_) by merging all the residues belonging to the three relevant sites mentioned above to avoid any bias towards any ligand-specific binding site.

To test the accuracy of our protocol in reproducing the conformation of different experimental binding sites of ADK, we selected four complex structures, each bearing a different ligand and displaying full sequence identity to that of the apo protein. The PDB IDs of these complexes are 1AKE,^92^ 2ECK,^91^ 1ANK,^54^ and 6F7U.^63^ In 1AKE, the protein was resolved in complex with the inhibitor AP5 (see **Figure 1**), mimicking the presence of two physiological substrates and binding across the LC and NC interfaces. In 1ANK, the protein was complexed with AMP, a physiological substrate, and with ANP, a non-hydrolysable ATP analog, while in the structure 2ECK the enzyme is bound to two physiological substrates, namely ADP and AMP. The non-hydrolysable (and thus catalytically incompetent) GTP analog GCP was bound to the LC interface of the protein in the 6F7U structure, which resembles closely the open (unbound) ADK structure. This latter system allows benchmarking the protocol against the impact of subtle chemical details on the conformation of the complexes and in turn on the structure-activity relationships for ADK. Four different experimental binding regions, labeled BS_AP5_, BS_GCP_, BS_ADP_, and BS_AMP_, were defined by taking all the residues within 3.5 Å from the corresponding ligands AP5, GCP, ADP, and AMP in their respective co-structures (**Table S1**). BS_AP5_ spans both the NC and LC interfaces, while BS_GCP_ and BS_ADP_ identify two slightly different regions across the LC interface, and BS_AMP_ is located within the NC interface. Note that, while enhanced conformational sampling was performed by biasing BS_COACH_, the sampling performance (that is the ability of gEDES in reproducing ADK bound-like conformations) was assessed with respect to the experimental binding regions, which are truly relevant to ligand binding.

### Standard MD

Standard all-atom MD simulations of the apo protein (hereafter MD_std_) embedded in a 0.15 KCl water solution (∼46.000 atoms in total) were carried out using the *pmemd* module of the AMBER20 package.^107^ The initial distance between the protein and the edge of the box was set to be at least 16 Å in each direction.

The topology file was created using the *LEaP* module of AmberTools21 starting from the apo structure. The AMBER-14SB^108^ force field was used for the protein, the TIP3P^109^ model was used for water, and the parameters for the ions were obtained from Wang *et al*.^110^ Long-range electrostatics was evaluated through the particle-mesh Ewald algorithm using a real-space cutoff of 12 Å and a grid spacing of 1 Å in each dimension. Van der Waals interactions were treated by a Lennard-Jones potential using a smooth cutoff (switching radius 10 Å, cutoff radius 12 Å). A multistep energy minimization with a combination of the steepest-descent and conjugate-gradient methods was carried out to relax the internal constraints of the systems by gradually releasing positional restraints. Next, the system was heated from 0 to 310 K in 1 ns of constant-pressure heating (NPT) MD simulation using the Langevin thermostat (collision frequency of 1 ps^-1^) and the Berendsen barostat. After equilibration (10 additional ns), four production runs of 2.5 μs each were performed, for a total of 10 μs. Time steps of 2 and 4 fs (after hydrogen mass repartitioning) were used respectively for pre-production and equilibrium NPT MD simulations. Coordinates from production trajectory were saved every 100 ps.

### Enhanced sampling MD

Bias-exchange well-tempered metadynamics simulations^43,44^ were performed on the apo protein using a set of *ad-hoc* collective variables (CVs) to enhance the sampling of both the shape and the volume of the binding pocket. We used the GROMACS 2022.4 package^111^ and the PLUMED 2.8 plugin.^112^ AMBER parameters were ported to GROMACS using the *acpype* parser.^113^ We defined four CVs considering the residues lining BS_COACH_: i) the radius of gyration (hereafter RoG_BS_) calculated using the *gyration* built-in function of PLUMED; ii) the number of (pseudo)contacts across three orthogonal “inertia planes” (CIPs), calculated through a switching function implemented in the *coordination* keyword of PLUMED. The “inertia planes” are defined as the planes orthogonal to the three principal inertia axes of the binding site and passing through its geometrical center. All non-hydrogenous atoms were considered to define the three CIPs, while only backbone atoms were used for RoG_BS_.

Starting from the last conformation sampled along the pre-production step in MD_std_, each replica was simulated without restraints for the first 10 ns. Next, an upper restraint centered at the value of RoG_X-ray_^apo^ was imposed with a force constant increasing linearly from 10 to 25 kcal mol^-1^Å^-2^ in 40 ns. This preliminary phase is needed to push the system towards a structure featuring a RoG_BS_ value close to RoG _X-ray_ ^apo^. Subsequently, the center of the RoG_BS_ restraint was decreased linearly (every ns) to a value corresponding to 85% of RoG_X-ray_ ^apo^ in 400 ns (moving restraints were applied on this CV -**Figure S2**). Focusing on conformations featuring collapsed sites is based on the evidence that the binding of ligands to enzymes is most often associated with such structural changes. Nonetheless, the relatively soft upper restraints adopted here still allow the sampling of structures with a larger RoG_BS_ with respect to the value at which the restraint is set (**Figure S2**). After reaching the desired value, the RoG_BS_ restraint remains active for 150 additional ns. Thus, the cumulative simulation time of each replica amounts to 550 ns.

To expand the use of our protocol for allosteric proteins bearing multiple binding sites, we employed SPECTRUS^114^ to identify the putative quasi-rigid domains of ADK. This analysis confirmed the dissection of the protein into three quasi-rigid domains, namely the CORE, LID, and NMP domains known from previous literature^74,114^ (**Figure 1**). The reliability of such subdivision was further verified by assessing, via RMSD calculations between the experimental structures, the internal conformational changes occurring upon ligand binding in each domain (**Table S2**). To account for hinge-like motions between the two LC and NC interfaces, we implemented the cRD CVs representing the number of (pseudo)contacts across these interfaces. To avoid relying on any experimental knowledge of the binding region, we only exploited the knowledge of BS_COACH_ to identify residues lining the NC and LC (sub)pockets. Namely, the following procedure was adopted: *i)* we selected all BS_COACH_ residues belonging to the NMP(LID) and being within 8 Å from any residue of the CORE; *ii)* a second specular selection was made by taking all BS_COACH_ residues belonging to the CORE that are within 8 Å from any residue of the NMP(LID). This cutoff was chosen to ensure that none of the BS_COACH_ residues got associated with more than one cRD CV and that each list contained a minimum of 4 residues belonging to each quasi-rigid domain (to limit the onset of large structural distortions within secondary structure elements). The union of selections *i)* and *ii)* defined the cumulative list of residues used to setup the cRD_LC(NC)_ CV (**Figure 2**); *iii)* the residues associated with the NC and LC (sub)pockets were split into two lists via domain assignment, for which the number of (pseudo)contacts was calculated via the *coordination* keyword of PLUMED. Furthermore, within each subpocket, the charged amino acids were separated from the others; this splitting in two CVs (cRD_NC(LC)c_ and cRD_NC(LC)o_) specifically enhances the conformational sampling of charged amino acids in targets, such as ADK, which contain many of them within the BS.

To facilitate the general applicability of the method, we automatized the workflow so that the latter cRD subdivision occurs only if: *i)* the binding site presents more than the 25% of charged residues and *ii)* each residue group defining a cRD variable is composed of at least 2 non-adjacent residues. For the system considered in this work, 11 out of the 32 residues (∼34 %) composing BS_COACH_ are charged, so this subdivision was applied.

The height *w* of the Gaussian hills was set to 0.6 kcal/mol, while their widths were set as reported in **Table S3** based on the fluctuations recorded during a short (∼200 ps) unbiased MD run. The bias factor for well-tempered metadynamics was set to 10. Hills were added every 2.5 ps, while the bias-exchange frequency was set to 50 ps. Further details on CV definitions are reported in **Table S3**. Coordinates of the system were saved every 10 ps.

### Cluster analysis

The cluster analysis was performed in the CVs space (for both the gEDES and MD_std_ simulations) using *in-house* scripts in the R language. The distribution of RoG_BS_ values sampled during the MD simulation was binned into 30 equally wide slices, and the built-in *hclust* module of R was used to perform a hierarchical agglomerative clustering within each slice, setting the number of generated clusters in the *i*^*th*^ slice to *x*_*i*_ *= (N*_*i*_*/N*_*tot*_*)·N*_*c*_, where *N*_*i*_, *N*_*tot*_, and *N*_*c*_ are respectively the number of structures within the *i*^*th*^ slice, the total number of structures, and the total number of clusters. In our case, *N*_*c*_ was set to 100, but we imposed the additional requirement to have at least two clusters within each of the 30 slices, which could thus lead to a final number of clusters larger than 100. This was implemented by iteratively increasing *N*_*c*_ by 10 units until the number of clusters within each of the RoG_BS_ slices was equal or higher than two. The resulting clusters were used as starting points to perform a second cluster analysis with the K-means method (maximum number of iterations set to 10000) and generating the same number of clusters. This multi-step strategy of clustering in the CV space, which outperforms more standard RMSD-based approaches in generating maximally diverse ensemble of protein conformations, resulted respectively in 130 and 160 clusters for the gEDES and MD_std_ trajectories.

### Docking

Docking calculations were performed using HADDOCK^93^ and AutoDock.^94^ Ligand conformations were extracted from the relative complex structures and prepared according to the standard procedure of each software. Docking calculations were performed for all ligands onto the predicted binding site (BS_COACH_); thus, ligands binding to the NC(LC) interface could in principle sample the LC(NC) one.

In HADDOCK, a single docking run was performed per case, starting from the various ensembles of *N*_*c*_ conformations, with increased sampling (50000/1000/1000 models for *it0, it1* and *wat* steps, respectively referring to rigid-body docking, semiflexible and final refinement in explicit solvent) using the HADDOCK2.4 web server.^115^ This increased sampling compared to the default was chosen to ensure that each conformation in the ensemble was sufficiently sampled. During *it0* the protein BS residues were defined as “active”, effectively drawing the rigid ligand into the BS without restraining its orientation. For the subsequent stages, only the ligand was active, improving its exploration of the binding site while maintaining at least one contact with its interacting residues. In addition, a fake bead was placed in the center of each binding site pocket and an ambiguous 0 Å distance restrain was defined to those two beads such as a ligand atom (any atom) should overlap with one of those two beads. Those two beads were defined as “shape” in the server and have no interactions with the remaining of the system except for the defined distance restraints. All conformations in the MD ensemble were aligned on the initial apo structure and their position, together with that of the fake beads, was kept fixed during the rigid-body phase of the docking.

In AutoDock, each ligand conformation was rigidly docked on each protein structure using the Lamarckian Genetic Algorithm (LGA). The number of energy evaluations (*ga_num_evals* parameter) was increased 10 times from its default value (1) to avoid repeating each calculation several times to obtain converged results. An adaptive grid was used, enclosing all the residues belonging to the BS in each different protein conformation.^39,40^ Finally, an additional step consisting in the relaxation of the docking poses by means of a restrained structural optimization was performed with AMBER20.^107^ Systems were relaxed *in vacuum* by means of up to 1000 cycles of steepest descent optimization followed by up to 24000 cycles using the conjugate gradients algorithm. Harmonic forces of 0.1 kcal·mol^-1^·Å^-1^ were applied on all non-hydrogenous atoms of the system. Long-range electrostatics was evaluated directly using a cutoff of 99 Å, as for the Lennard-Jones potential. The AMBER-14SB^108^ force field was used for the protein, while the parameters of the ligands were derived from the GAFF^116^ force field using the *antechamber* module of AmberTools. Bond-charge corrections (bcc) charges were assigned to ligand atoms following structural relaxation under the “Austin Model 1 (AM1)” approximation. After this step, poses were scored according to AutoDock’s energy function. Next, the top poses (in total *N*_*c*_, one for each docking run performed on a different receptor structure) were clustered using the *cpptraj* module of AmberTools with a hierarchical agglomerative algorithm. Namely, after structural alignment of the BS for the different complex conformations, ligand poses were clustered using a distance RMSD (dRMSD) cutoff *d*_*c*_ = *0*.*067* · *N*_*nh*_, where *N*_*nh*_ is the number of non-hydrogenous atoms of the ligand. This choice was made to tune the cutoff to the molecular size of each compound and resulted in cutoffs of 3.8, 1.5, 1.8, and 2.1 Å for AP5, AMP, ADP, and GCP, respectively. Finally, clusters were ordered according to the top score (lowest binding free energy) within each cluster.

### Complex structure prediction with NeuralPLexer and AlphaFold3

NeuralPLexer^32^ simulations were run giving in input as *receptor* and *template* the chain A of the apo-structure of ADK. In addition, for each ligand, the molecular structure extracted from the PDB file of the corresponding complex was given as input to the *ligand* keyword. All files were provided in pdb format, and the remaining parameters were set to their default values (*num-steps = 40, sampler = Langevin simulated annealing*). The *‘batched structure sampling’* method was employed, which produced 16 (keyword *n-samples*) putative structures of the corresponding protein-ligand complex. The *‘complex structure prediction’* model, specifically trained for complex structure prediction, was used as model checkpoint.

AlphaFold3^33^ predictions were performed on the website https://alphafoldserver.com by selecting ADP or AMP as ligand and inserting the FASTA sequence of ADK corresponding to the PDB structure with PDB ID 4AKE.

## Acknowledgments

This project was funded by the Regional Government of Sardinia through the European Regional Development Fund grant “PROOF of CONCEPT - Valorizzazione dei risultati della ricerca in biomedicina”. M.A. and A.V.V. gratefully acknowledge the “One Health Basic and Translational Research Actions addressing Unmet Needs on Emerging Infectious Diseases (INF-ACT)” foundation by the Italian Ministry of University and Research, PNRR, mission 4, component 2, investment 1.3, project number PE00000007 (University of Cagliari). G.M., A. Bosin and P.R. gratefully acknowledge the Health Extended Alliance for Innovative Therapies, Advanced Lab-research, and Integrated Approaches of Precision Medicine partnership (HEAL ITALIA), foundation by the Italian Ministry of University and Research, PNRR, mission 4, component 2, investment 1.3, project number PE00000019 (University of Cagliari). G.M, A. Bosin, P.R. and A.V.V. acknowledge funding from the National Recovery and Resilience Plan (NRRP), Mission 4 Component 2 Investment 1.5 - Call for tender No.3277 published on December 30, 2021 by the Italian Ministry of University and Research (MUR) funded by the European Union – NextGenerationEU. Project Code ECS0000038 – Project Title eINS Ecosystem of Innovation for Next Generation Sardinia – CUP J85B17000360007 - Concession Decree No. 1056 adopted on June 23, 2022 by the Italian Ministry of University and Research (MUR). A.B., M. A., G.M., P.R., and A.V.V. received financial support by the NIAID/NIH grant no. R01AI136799. A.M.J.J.B. acknowledges funding from the European Union Horizon 2020 project BioExcel (823830). We thank Vincenzo Carnevale (Temple University, Philadelphia, U.S.A.) for useful discussions and Mattia Bernetti (Istituto Italiano di Tecnologia, Italy) and Paolo Carloni (Forschungszentrum Jülich GmbH, Germany) for the careful reading of the manuscript.

## Patent Application

A patent application for the gEDES protocol has been filed with the Italian Patent and Trademark Office (Ufficio Italiano Brevetti e Marchi) under the application number 102023000013206.

## References

(1) Du, X.; Li, Y.; Xia, Y.-L.; Ai, S.-M.; Liang, J.; Sang, P.; Ji, X.-L.; Liu, S.-Q. Insights into Protein–Ligand Interactions: Mechanisms, Models, and Methods. IJMS 2016, 17 (2), 144. 10.3390/ijms17020144.

(2) Chu, W.-T.; Yan, Z.; Chu, X.; Zheng, X.; Liu, Z.; Xu, L.; Zhang, K.; Wang, J. Physics of Biomolecular Recognition and Conformational Dynamics. Rep. Prog. Phys. 2021, 84 (12), 126601. 10.1088/1361-6633/ac3800.

(3) Ma, B.; Shatsky, M.; Wolfson, H. J.; Nussinov, R. Multiple Diverse Ligands Binding at a Single Protein Site: A Matter of Pre-existing Populations. Protein Science 2002, 11 (2), 184–197. 10.1110/ps.21302.

(4) Niitsu, A.; Re, S.; Oshima, H.; Kamiya, M.; Sugita, Y. De Novo Prediction of Binders and Nonbinders for T4 Lysozyme by gREST Simulations. J. Chem. Inf. Model. 2019, 59 (9), 3879–3888. 10.1021/acs.jcim.9b00416.

(5) Ahuja, L. G.; Aoto, P. C.; Kornev, A. P.; Veglia, G.; Taylor, S. S. Dynamic Allostery-Based Molecular Workings of Kinase:Peptide Complexes. Proc. Natl. Acad. Sci. U.S.A. 2019, 116 (30), 15052–15061. 10.1073/pnas.1900163116.

(6) Bonomi, M.; Camilloni, C.; Cavalli, A.; Vendruscolo, M. Metainference: A Bayesian Inference Method for Heterogeneous Systems. Sci. Adv. 2016, 2 (1), e1501177. 10.1126/sciadv.1501177.

(7) Bottaro, S.; Lindorff-Larsen, K. Biophysical Experiments and Biomolecular Simulations: A Perfect Match? Science 2018, 361 (6400), 355–360. 10.1126/science.aat4010.

(8) Senior, A. W.; Evans, R.; Jumper, J.; Kirkpatrick, J.; Sifre, L.; Green, T.; Qin, C.; Žídek, A.; Nelson, A. W. R.; Bridgland, A.; Penedones, H.; Petersen, S.; Simonyan, K.; Crossan, S.; Kohli, P.; Jones, D. T.; Silver, D.; Kavukcuoglu, K.; Hassabis, D. Improved Protein Structure Prediction Using Potentials from Deep Learning. Nature 2020, 577 (7792), 706–710. 10.1038/s41586-019-1923-7.

(9) Ganesan, A.; Coote, M. L.; Barakat, K. Molecular Dynamics-Driven Drug Discovery: Leaping Forward with Confidence. Drug Discovery Today 2017, 22 (2), 249–269. 10.1016/j.drudis.2016.11.001.

(10) Śledź, P.; Caflisch, A. Protein Structure-Based Drug Design: From Docking to Molecular Dynamics. Current Opinion in Structural Biology 2018, 48, 93–102. 10.1016/j.sbi.2017.10.010.

(11) Allison, J. R. Computational Methods for Exploring Protein Conformations. Biochemical Society Transactions 2020, 48 (4), 1707–1724. 10.1042/BST20200193.

(12) De Vivo, M.; Masetti, M.; Bottegoni, G.; Cavalli, A. Role of Molecular Dynamics and Related Methods in Drug Discovery. J. Med. Chem. 2016, 59 (9), 4035–4061. 10.1021/acs.jmedchem.5b01684.

(13) Muscat, S.; Stojceski, F.; Danani, A. Elucidating the Effect of Static Electric Field on Amyloid Beta 1–42 Supramolecular Assembly. Journal of Molecular Graphics and Modelling 2020, 96, 107535. 10.1016/j.jmgm.2020.107535.

(14) Lazim, R.; Suh, D.; Choi, S. Advances in Molecular Dynamics Simulations and Enhanced Sampling Methods for the Study of Protein Systems. IJMS 2020, 21 (17), 6339. 10.3390/ijms21176339.

(15) Bouvier, B. Curvature as a Collective Coordinate in Enhanced Sampling Membrane Simulations. J. Chem. Theory Comput. 2019, 15 (12), 6551–6561. 10.1021/acs.jctc.9b00716.

(16) Souza, P. C. T.; Thallmair, S.; Conflitti, P.; Ramírez-Palacios, C.; Alessandri, R.; Raniolo, S.; Limongelli, V.; Marrink, S. J. Protein–Ligand Binding with the Coarse-Grained Martini Model. Nat Commun 2020, 11 (1), 3714. 10.1038/s41467-020-17437-5.

(17) Heilmann, N.; Wolf, M.; Kozlowska, M.; Sedghamiz, E.; Setzler, J.; Brieg, M.; Wenzel, W. Sampling of the Conformational Landscape of Small Proteins with Monte Carlo Methods. Sci Rep 2020, 10 (1), 18211. 10.1038/s41598-020-75239-7.

(18) Sasmal, S.; Gill, S. C.; Lim, N. M.; Mobley, D. L. Sampling Conformational Changes of Bound Ligands Using Nonequilibrium Candidate Monte Carlo and Molecular Dynamics. J. Chem. Theory Comput. 2020, 16 (3), 1854–1865. 10.1021/acs.jctc.9b01066.

(19) Degiacomi, M. T. Coupling Molecular Dynamics and Deep Learning to Mine Protein Conformational Space. Structure 2019, 27 (6), 1034-1040.e3. 10.1016/j.str.2019.03.018.

(20) Noé, F.; De Fabritiis, G.; Clementi, C. Machine Learning for Protein Folding and Dynamics. Current Opinion in Structural Biology 2020, 60, 77–84. 10.1016/j.sbi.2019.12.005.

(21) Yuan, J.-H.; Han, S. B.; Richter, S.; Wade, R. C.; Kokh, D. B. Druggability Assessment in TRAPP Using Machine Learning Approaches. J. Chem. Inf. Model. 2020, 60 (3), 1685–1699. 10.1021/acs.jcim.9b01185.

(22) Jumper, J.; Evans, R.; Pritzel, A.; Green, T.; Figurnov, M.; Ronneberger, O.; Tunyasuvunakool, K.; Bates, R.; Žídek, A.; Potapenko, A.; Bridgland, A.; Meyer, C.; Kohl, S. A. A.; Ballard, A. J.; Cowie, A.; Romera-Paredes, B.; Nikolov, S.; Jain, R.; Adler, J.; Back, T.; Petersen, S.; Reiman, D.; Clancy, E.; Zielinski, M.; Steinegger, M.; Pacholska, M.; Berghammer, T.; Bodenstein, S.; Silver, D.; Vinyals, O.; Senior, A. W.; Kavukcuoglu, K.; Kohli, P.; Hassabis, D. Highly Accurate Protein Structure Prediction with AlphaFold. Nature 2021, 596 (7873), 583–589. 10.1038/s41586-021-03819-2.

(23) Motta, S.; Callea, L.; Bonati, L.; Pandini, A. PathDetect-SOM: A Neural Network Approach for the Identification of Pathways in Ligand Binding Simulations. J. Chem. Theory Comput. 2022, 18 (3), 1957–1968. 10.1021/acs.jctc.1c01163.

(24) Zhang, Y.; Li, S.; Meng, K.; Sun, S. Machine Learning for Sequence and Structure-Based Protein–Ligand Interaction Prediction. J. Chem. Inf. Model. 2024, 64 (5), 1456–1472. 10.1021/acs.jcim.3c01841.

(25) Müllender, L.; Rizzi, A.; Parrinello, M.; Carloni, P.; Mandelli, D. Effective Data-Driven Collective Variables for Free Energy Calculations from Metadynamics of Paths. PNAS Nexus 2024, 3 (4), pgae159. 10.1093/pnasnexus/pgae159.

(26) Amaro, R. E.; Baudry, J.; Chodera, J.; Demir, Ö.; McCammon, J. A.; Miao, Y.; Smith, J. C. Ensemble Docking in Drug Discovery. Biophysical Journal 2018, 114 (10), 2271–2278. 10.1016/j.bpj.2018.02.038.

(27) Buonfiglio, R.; Recanatini, M.; Masetti, M. Protein Flexibility in Drug Discovery: From Theory to Computation. ChemMedChem 2015, 10 (7), 1141–1148. 10.1002/cmdc.201500086.

(28) Antunes, D. A.; Devaurs, D.; Kavraki, L. E. Understanding the Challenges of Protein Flexibility in Drug Design. Expert Opinion on Drug Discovery 2015, 10 (12), 1301–1313. 10.1517/17460441.2015.1094458.

(29) Caballero, J. The Latest Automated Docking Technologies for Novel Drug Discovery. Expert Opinion on Drug Discovery 2020, 1–21. 10.1080/17460441.2021.1858793.

(30) Harmalkar, A.; Gray, J. J. Advances to Tackle Backbone Flexibility in Protein Docking. Current Opinion in Structural Biology 2021, 67, 178–186. 10.1016/j.sbi.2020.11.011.

(31) Saldaño, T.; Escobedo, N.; Marchetti, J.; Zea, D. J.; Mac Donagh, J.; Velez Rueda, A.J.; Gonik, E.; García Melani, A.; Novomisky Nechcoff, J.; Salas, M. N.; Peters, T.; Demitroff, N.; Fernandez Alberti, S.; Palopoli, N.; Fornasari, M. S.; Parisi, G. Impact of Protein Conformational Diversity on AlphaFold Predictions. Bioinformatics 2022, 38 (10), 2742–2748. 10.1093/bioinformatics/btac202.

(32) Qiao, Z.; Nie, W.; Vahdat, A.; Miller, T. F.; Anandkumar, A. State-Specific Protein–Ligand Complex Structure Prediction with a Multiscale Deep Generative Model. Nat Mach Intell 2024, 6 (2), 195–208. 10.1038/s42256-024-00792-z.

(33) Abramson, J.; Adler, J.; Dunger, J.; Evans, R.; Green, T.; Pritzel, A.; Ronneberger, O.; Willmore, L.; Ballard, A. J.; Bambrick, J.; Bodenstein, S. W.; Evans, D. A.; Hung, C.-C.; O’Neill, M.; Reiman, D.; Tunyasuvunakool, K.; Wu, Z.; Žemgulytė, A.; Arvaniti, E.; Beattie, C.; Bertolli, O.; Bridgland, A.; Cherepanov, A.; Congreve, M.; Cowen-Rivers, A. I.; Cowie, A.; Figurnov, M.; Fuchs, F. B.; Gladman, H.; Jain, R.; Khan, Y. A.; Low, C. M. R.; Perlin, K.; Potapenko, A.; Savy, P.; Singh, S.; Stecula, A.; Thillaisundaram, A.; Tong, C.; Yakneen, S.; Zhong, E. D.; Zielinski, M.; Žídek, A.; Bapst, V.; Kohli, P.; Jaderberg, M.; Hassabis, D.; Jumper, J. M. Accurate Structure Prediction of Biomolecular Interactions with AlphaFold 3. Nature 2024, 630 (8016), 493–500. 10.1038/s41586-024-07487-w.

(34) Huang, S.-Y.; Zou, X. Ensemble Docking of Multiple Protein Structures: Considering Protein Structural Variations in Molecular Docking. Proteins 2006, 66 (2), 399–421. 10.1002/prot.21214.

(35) Kapoor, K.; Thangapandian, S.; Tajkhorshid, E. Extended-Ensemble Docking to Probe Dynamic Variation of Ligand Binding Sites during Large-Scale Structural Changes of Proteins. Chem. Sci. 2022, 13 (14), 4150–4169. 10.1039/D2SC00841F.

(36) Acharya, A.; Agarwal, R.; Baker, M. B.; Baudry, J.; Bhowmik, D.; Boehm, S.; Byler, K. G.; Chen, S. Y.; Coates, L.; Cooper, C. J.; Demerdash, O.; Daidone, I.; Eblen, J. D.; Ellingson, S.; Forli, S.; Glaser, J.; Gumbart, J. C.; Gunnels, J.; Hernandez, O.; Irle, S.; Kneller, D. W.; Kovalevsky, A.; Larkin, J.; Lawrence, T. J.; LeGrand, S.; Liu, S.-H.; Mitchell, J. C.; Park, G.; Parks, J. M.; Pavlova, A.; Petridis, L.; Poole, D.; Pouchard, L.; Ramanathan, A.; Rogers, D. M.; Santos-Martins, D.; Scheinberg, A.; Sedova, A.; Shen, Y.; Smith, J. C.; Smith, M. D.; Soto, C.; Tsaris, A.; Thavappiragasam, M.; Tillack, A. F.; Vermaas, J. V.; Vuong, V. Q.; Yin, J.; Yoo, S.; Zahran, M.; Zanetti-Polzi, L. Supercomputer-Based Ensemble Docking Drug Discovery Pipeline with Application to Covid-19. J. Chem. Inf. Model. 2020, 60 (12), 5832–5852. 10.1021/acs.jcim.0c01010.

(37) Akbari, Z.; Stagno, C.; Iraci, N.; Efferth, T.; Omer, E. A.; Piperno, A.; Montazerozohori, M.; Feizi-Dehnayebi, M.; Micale, N. Biological Evaluation, DFT, MEP, HOMO-LUMO Analysis and Ensemble Docking Studies of Zn(II) Complexes of Bidentate and Tetradentate Schiff Base Ligands as Antileukemia Agents. Journal of Molecular Structure 2024, 1301, 137400. 10.1016/j.molstruc.2023.137400.

(38) Ricci-Lopez, J.; Aguila, S. A.; Gilson, M. K.; Brizuela, C. A. Improving Structure-Based Virtual Screening with Ensemble Docking and Machine Learning. J. Chem. Inf. Model. 2021, 61 (11), 5362–5376. 10.1021/acs.jcim.1c00511.

(39) Basciu, A.; Malloci, G.; Pietrucci, F.; Bonvin, A. M. J. J.; Vargiu, A. V. Holo-like and Druggable Protein Conformations from Enhanced Sampling of Binding Pocket Volume and Shape. J. Chem. Inf. Model. 2019, 59 (4), 1515–1528. 10.1021/acs.jcim.8b00730.

(40) Basciu, A.; Koukos, P. I.; Malloci, G.; Bonvin, A. M. J. J.; Vargiu, A. V. Coupling Enhanced Sampling of the Apo-Receptor with Template-Based Ligand Conformers Selection: Performance in Pose Prediction in the D3R Grand Challenge 4. J Comput Aided Mol Des 2020, 34 (2), 149–162. 10.1007/s10822-019-00244-6.

(41) Basciu, A.; Callea, L.; Motta, S.; Bonvin, A. M. J. J.; Bonati, L.; Vargiu, A. V. No Dance, No Partner! A Tale of Receptor Flexibility in Docking and Virtual Screening. In Annual Reports in Medicinal Chemistry; Elsevier, 2022; Vol. 59, pp 43–97. 10.1016/bs.armc.2022.08.006.

(42) Laio, A.; Parrinello, M. Escaping Free-Energy Minima. Proceedings of the National Academy of Sciences 2002, 99 (20), 12562–12566. 10.1073/pnas.202427399.

(43) Barducci, A.; Bussi, G.; Parrinello, M. Well-Tempered Metadynamics: A Smoothly Converging and Tunable Free-Energy Method. Phys. Rev. Lett. 2008, 100 (2), 020603. 10.1103/PhysRevLett.100.020603.

(44) Piana, S.; Laio, A. A Bias-Exchange Approach to Protein Folding. J. Phys. Chem. B 2007, 111 (17), 4553–4559. 10.1021/jp067873l.

(45) Zuo, K.; Kranjc, A.; Capelli, R.; Rossetti, G.; Nechushtai, R.; Carloni, P. Metadynamics Simulations of Ligands Binding to Protein Surfaces: A Novel Tool for Rational Drug Design. Phys. Chem. Chem. Phys. 2023, 25 (20), 13819–13824. 10.1039/D3CP01388J.

(46) Cavalli, A.; Spitaleri, A.; Saladino, G.; Gervasio, F. L. Investigating Drug–Target Association and Dissociation Mechanisms Using Metadynamics-Based Algorithms. Acc. Chem. Res. 2015, 48 (2), 277–285. 10.1021/ar500356n.

(47) Nussinov, R.; Tsai, C.-J. Allostery in Disease and in Drug Discovery. Cell 2013, 153 (2), 293–305. 10.1016/j.cell.2013.03.034.

(48) Sheik Amamuddy, O.; Veldman, W.; Manyumwa, C.; Khairallah, A.; Agajanian, S.; Oluyemi, O.; Verkhivker, G. M.; Tastan Bishop, Ö. Integrated Computational Approaches and Tools for Allosteric Drug Discovery. IJMS 2020, 21 (3), 847. 10.3390/ijms21030847.

(49) Ionescu, M. I. Adenylate Kinase: A Ubiquitous Enzyme Correlated with Medical Conditions. Protein J 2019, 38 (2), 120–133. 10.1007/s10930-019-09811-0.

(50) Carling, D. AMPK Signalling in Health and Disease. Current Opinion in Cell Biology 2017, 45, 31–37. 10.1016/j.ceb.2017.01.005.

(51) Müller, C.; Schlauderer, G.; Reinstein, J.; Schulz, G. Adenylate Kinase Motions during Catalysis: An Energetic Counterweight Balancing Substrate Binding. Structure 1996, 4 (2), 147–156. 10.1016/S0969-2126(96)00018-4.

(52) Bae, E.; Phillips, G. N. Roles of Static and Dynamic Domains in Stability and Catalysis of Adenylate Kinase. PNAS 2006, 103 (7), 2132–2137. 10.1073/pnas.0507527103.

(53) Liu, R.; Xu, H.; Wei, Z.; Wang, Y.; Lin, Y.; Gong, W. Crystal Structure of Human Adenylate Kinase 4 (L171P) Suggests the Role of Hinge Region in Protein Domain Motion. Biochemical and Biophysical Research Communications 2009, 379 (1), 92–97. 10.1016/j.bbrc.2008.12.012.

(54) Berry, M. B.; Meador, B.; Bilderback, T.; Liang, P.; Glaser, M.; Phillips, G. N. The Closed Conformation of a Highly Flexible Protein: The Structure ofE. Coli Adenylate Kinase with Bound AMP and AMPPNP. Proteins 1994, 19 (3), 183–198. 10.1002/prot.340190304.

(55) Schulz, G. E.; Müller, C. W.; Diederichs, K. Induced-Fit Movements in Adenylate Kinases. Journal of Molecular Biology 1990, 213 (4), 627–630. 10.1016/S0022-2836(05)80250-5.

(56) Wu, P.; Nielsen, T. E.; Clausen, M. H. FDA-Approved Small-Molecule Kinase Inhibitors. Trends in Pharmacological Sciences 2015, 36 (7), 422–439. 10.1016/j.tips.2015.04.005.

(57) Arrowsmith, C. H.; Bountra, C.; Fish, P. V.; Lee, K.; Schapira, M. Epigenetic Protein Families: A New Frontier for Drug Discovery. Nat Rev Drug Discov 2012, 11 (5), 384–400. 10.1038/nrd3674.

(58) Whitford, P. C.; Gosavi, S.; Onuchic, J. N. Conformational Transitions in Adenylate Kinase: ALLOSTERIC COMMUNICATION REDUCES MISLIGATION. J. Biol. Chem. 2008, 283 (4), 2042–2048. 10.1074/jbc.M707632200.

(59) Potoyan, D. A.; Zhuravlev, P. I.; Papoian, G. A. Computing Free Energy of a Large-Scale Allosteric Transition in Adenylate Kinase Using All Atom Explicit Solvent Simulations. J. Phys. Chem. B 2012, 116 (5), 1709–1715. 10.1021/jp209980b.

(60) Daily, M. D.; Phillips, G. N.; Cui, Q. Many Local Motions Cooperate to Produce the Adenylate Kinase Conformational Transition. Journal of Molecular Biology 2010, 400 (3), 618–631. 10.1016/j.jmb.2010.05.015.

(61) Kerns, S. J.; Agafonov, R. V.; Cho, Y.-J.; Pontiggia, F.; Otten, R.; Pachov, D. V.; Kutter, S.; Phung, L. A.; Murphy, P. N.; Thai, V.; Alber, T.; Hagan, M. F.; Kern, D. The Energy Landscape of Adenylate Kinase during Catalysis. Nat Struct Mol Biol 2015, 22 (2), 124–131. 10.1038/nsmb.2941.

(62) Lassila, J. K.; Zalatan, J. G.; Herschlag, D. Biological Phosphoryl-Transfer Reactions: Understanding Mechanism and Catalysis. Annu. Rev. Biochem. 2011, 80 (1), 669–702. 10.1146/annurev-biochem-060409-092741.

(63) Rogne, P.; Rosselin, M.; Grundström, C.; Hedberg, C.; Sauer, U. H.; Wolf-Watz, M. Molecular Mechanism of ATP versus GTP Selectivity of Adenylate Kinase. Proc Natl Acad Sci USA 2018, 115 (12), 3012–3017. 10.1073/pnas.1721508115.

(64) Vonrhein, C.; Schlauderer, G. J.; Schulz, G. E. Movie of the Structural Changes during a Catalytic Cycle of Nucleoside Monophosphate Kinases. Structure 1995, 3 (5), 483–490. 10.1016/S0969-2126(01)00181-2.

(65) Kovermann, M.; Grundström, C.; Sauer-Eriksson, A. E.; Sauer, U. H.; Wolf-Watz, M. Structural Basis for Ligand Binding to an Enzyme by a Conformational Selection Pathway. Proc Natl Acad Sci USA 2017, 114 (24), 6298–6303. 10.1073/pnas.1700919114.

(66) Hanson, J. A.; Duderstadt, K.; Watkins, L. P.; Bhattacharyya, S.; Brokaw, J.; Chu, J.-W.; Yang, H. Illuminating the Mechanistic Roles of Enzyme Conformational Dynamics. Proceedings of the National Academy of Sciences 2007, 104 (46), 18055–18060. 10.1073/pnas.0708600104.

(67) Scheerer, D.; Adkar, B. V.; Bhattacharyya, S.; Levy, D.; Iljina, M.; Riven, I.; Dym, O.; Haran, G.; Shakhnovich, E. I. Allosteric Communication between Ligand Binding Domains Modulates Substrate Inhibition in Adenylate Kinase. Proc. Natl. Acad. Sci. U.S.A. 2023, 120 (18), e2219855120. 10.1073/pnas.2219855120.

(68) Ionescu, M.; Oniga, O. Molecular Docking Evaluation of (E)-5-Arylidene-2-Thioxothiazolidin-4-One Derivatives as Selective Bacterial Adenylate Kinase Inhibitors. Molecules 2018, 23 (5), 1076. 10.3390/molecules23051076.

(69) Li, D.; Liu, M. S.; Ji, B. Mapping the Dynamics Landscape of Conformational Transitions in Enzyme: The Adenylate Kinase Case. Biophysical Journal 2015, 109 (3), 647–660. 10.1016/j.bpj.2015.06.059.

(70) Kruger, D. M.; Ahmed, A.; Gohlke, H. NMSim Web Server: Integrated Approach for Normal Mode-Based Geometric Simulations of Biologically Relevant Conformational Transitions in Proteins. Nucleic Acids Research 2012, 40 (W1), W310–W316. 10.1093/nar/gks478.

(71) Ahmed, A.; Rippmann, F.; Barnickel, G.; Gohlke, H. A Normal Mode-Based Geometric Simulation Approach for Exploring Biologically Relevant Conformational Transitions in Proteins. Journal of Chemical Information and Modeling 2011, 51 (7), 1604–1622. 10.1021/ci100461k.

(72) Lu, Q.; Wang, J. Single Molecule Conformational Dynamics of Adenylate Kinase: Energy Landscape, Structural Correlations, and Transition State Ensembles. J. Am. Chem. Soc. 2008, 130 (14), 4772–4783. 10.1021/ja0780481.

(73) Seyler, S. L.; Beckstein, O. Sampling Large Conformational Transitions: Adenylate Kinase as a Testing Ground. Molecular Simulation 2014, 40 (10–11), 855–877. 10.1080/08927022.2014.919497.

(74) Formoso, E.; Limongelli, V.; Parrinello, M. Energetics and Structural Characterization of the Large-Scale Functional Motion of Adenylate Kinase. Sci Rep 2015, 5 (1), 8425. 10.1038/srep08425.

(75) Yasuda, T.; Shigeta, Y.; Harada, R. Efficient Conformational Sampling of Collective Motions of Proteins with Principal Component Analysis-Based Parallel Cascade Selection Molecular Dynamics. J. Chem. Inf. Model. 2020, 60 (8), 4021–4029. 10.1021/acs.jcim.0c00580.

(76) Jalalypour, F.; Sensoy, O.; Atilgan, C. Perturb–Scan–Pull: A Novel Method Facilitating Conformational Transitions in Proteins. J. Chem. Theory Comput. 2020, 16 (6), 3825–3841. 10.1021/acs.jctc.9b01222.

(77) Wang, J.; Peng, C.; Yu, Y.; Chen, Z.; Xu, Z.; Cai, T.; Shao, Q.; Shi, J.; Zhu, W. Exploring Conformational Change of Adenylate Kinase by Replica Exchange Molecular Dynamic Simulation. Biophysical Journal 2020, 118 (5), 1009–1018. 10.1016/j.bpj.2020.01.001.

(78) Uyar, A.; Kantarci-Carsibasi, N.; Haliloglu, T.; Doruker, P. Features of Large Hinge-Bending Conformational Transitions. Prediction of Closed Structure from Open State. Biophysical Journal 2014, 106 (12), 2656–2666. 10.1016/j.bpj.2014.05.017.

(79) Zeller, F.; Zacharias, M. Substrate Binding Specifically Modulates Domain Arrangements in Adenylate Kinase. Biophysical Journal 2015, 109 (9), 1978–1985. 10.1016/j.bpj.2015.08.049.

(80) Kurkcuoglu, Z.; Doruker, P. Ligand Docking to Intermediate and Close-To-Bound Conformers Generated by an Elastic Network Model Based Algorithm for Highly Flexible Proteins. PLoS ONE 2016, 11 (6), e0158063. 10.1371/journal.pone.0158063.

(81) Lou, H.; Cukier, R. I. Molecular Dynamics of Apo-Adenylate Kinase: A Principal Component Analysis. J. Phys. Chem. B 2006, 110 (25), 12796–12808. 10.1021/jp061976m.

(82) Nam, K.; Arattu Thodika, A. R.; Grundström, C.; Sauer, U. H.; Wolf-Watz, M. Elucidating Dynamics of Adenylate Kinase from Enzyme Opening to Ligand Release. J. Chem. Inf. Model. 2024, 64 (1), 150–163. 10.1021/acs.jcim.3c01618.

(83) Fiorentini, R.; Tarenzi, T.; Potestio, R. Fast, Accurate, and System-Specific Variable-Resolution Modeling of Proteins. J. Chem. Inf. Model. 2023, 63 (4), 1260–1275. 10.1021/acs.jcim.2c01311.

(84) Kanada, R.; Terayama, K.; Tokuhisa, A.; Matsumoto, S.; Okuno, Y. Enhanced Conformational Sampling with an Adaptive Coarse-Grained Elastic Network Model Using Short-Time All-Atom Molecular Dynamics. J. Chem. Theory Comput. 2022, 18 (4), 2062–2074. 10.1021/acs.jctc.1c01074.

(85) Yuan, Y.; Zhu, Q.; Song, R.; Ma, J.; Dong, H. A Two-Ended Data-Driven Accelerated Sampling Method for Exploring the Transition Pathways between Two Known States of Protein. J. Chem. Theory Comput. 2020, 16 (7), 4631–4640. 10.1021/acs.jctc.9b01184.

(86) Yang, S.; Song, C. Switching Go — -Martini for Investigating Protein Conformational Transitions and Associated Protein–Lipid Interactions. J. Chem. Theory Comput. 2024, 20 (6), 2618–2629. 10.1021/acs.jctc.3c01222.

(87) Wang, Y.; Makowski, L. Fine Structure of Conformational Ensembles in Adenylate Kinase. Proteins 2018, 86 (3), 332–343. 10.1002/prot.25443.

(88) Henzler-Wildman, K. A.; Thai, V.; Lei, M.; Ott, M.; Wolf-Watz, M.; Fenn, T.; Pozharski, E.; Wilson, M. A.; Petsko, G. A.; Karplus, M.; Hübner, C. G.; Kern, D. Intrinsic Motions along an Enzymatic Reaction Trajectory. Nature 2007, 450 (7171), 838–844. 10.1038/nature06410.

(89) Shao, Q. Enhanced Conformational Sampling Technique Provides an Energy Landscape View of Large-Scale Protein Conformational Transitions. Phys. Chem. Chem. Phys. 2016, 18 (42), 29170–29182. 10.1039/C6CP05634B.

(90) Punia, R.; Goel, G. Computation of the Protein Conformational Transition Pathway on Ligand Binding by Linear Response-Driven Molecular Dynamics. J. Chem. Theory Comput. 2022, 18 (5), 3268–3283. 10.1021/acs.jctc.1c01243.

(91) Berry, M. B.; Bae, E.; Bilderback, T. R.; Glaser, M.; Phillips, G. N. Crystal Structure of ADP/AMP Complex of Escherichia Coli Adenylate Kinase. Proteins 2005, 62 (2), 555–556. 10.1002/prot.20699.

(92) Müller, C. W.; Schulz, G. E. Structure of the Complex between Adenylate Kinase from Escherichia Coli and the Inhibitor Ap5A Refined at 1.9 Å Resolution. Journal of Molecular Biology 1992, 224 (1), 159–177. 10.1016/0022-2836(92)90582-5.

(93) Dominguez, C.; Boelens, R.; Bonvin, A. M. J. J. HADDOCK: A Protein−Protein Docking Approach Based on Biochemical or Biophysical Information. J. Am. Chem. Soc. 2003, 125 (7), 1731–1737. 10.1021/ja026939x.

(94) Morris, G. M.; Huey, R.; Lindstrom, W.; Sanner, M. F.; Belew, R. K.; Goodsell, D. S.; Olson, A. J. AutoDock4 and AutoDockTools4: Automated Docking with Selective Receptor Flexibility. J. Comput. Chem. 2009, 30 (16), 2785–2791. 10.1002/jcc.21256.

(95) Flores, S. C.; Gerstein, M. B. Predicting Protein Ligand Binding Motions with the Conformation Explorer. BMC Bioinformatics 2011, 12 (1), 417. 10.1186/1471-2105-12-417.

(96) Wang, J.; Peng, C.; Yu, Y.; Chen, Z.; Xu, Z.; Cai, T.; Shao, Q.; Shi, J.; Zhu, W. Exploring Conformational Change of Adenylate Kinase by Replica Exchange Molecular Dynamic Simulation. Biophysical Journal 2020, 118 (5), 1009–1018. 10.1016/j.bpj.2020.01.001.

(97) Amaro, R.; Li, W. Emerging Methods for Ensemble-Based Virtual Screening. CTMC 2010, 10 (1), 3–13. 10.2174/156802610790232279.

(98) Tarcsay, Á.; Paragi, G.; Vass, M.; Jójárt, B.; Bogár, F.; Keserű, G. M. The Impact of Molecular Dynamics Sampling on the Performance of Virtual Screening against GPCRs. J. Chem. Inf. Model. 2013, 53 (11), 2990–2999. 10.1021/ci400087b.

(99) Ferreira, L.; dos Santos, R.; Oliva, G.; Andricopulo, A. Molecular Docking and Structure-Based Drug Design Strategies. Molecules 2015, 20 (7), 13384–13421. 10.3390/molecules200713384.

(100) Forli, S. Charting a Path to Success in Virtual Screening. Molecules 2015, 20 (10), 18732–18758. 10.3390/molecules201018732.

(101) Li, Y. Y.; An, J.; Jones, S. J. M. A Computational Approach to Finding Novel Targets for Existing Drugs. PLoS Comput Biol 2011, 7 (9), e1002139. 10.1371/journal.pcbi.1002139.

(102) Schmidtke, P.; Barril, X. Understanding and Predicting Druggability. A High-Throughput Method for Detection of Drug Binding Sites. J. Med. Chem. 2010, 53 (15), 5858–5867. 10.1021/jm100574m.

(103) Miao, Y.; Goldfeld, D. A.; Moo, E. V.; Sexton, P. M.; Christopoulos, A.; McCammon, J. A.; Valant, C. Accelerated Structure-Based Design of Chemically Diverse Allosteric Modulators of a Muscarinic G Protein-Coupled Receptor. Proc Natl Acad Sci USA 2016, 113 (38), E5675–E5684. 10.1073/pnas.1612353113.

(104) Pandini, A.; Fornili, A. Using Local States To Drive the Sampling of Global Conformations in Proteins. J. Chem. Theory Comput. 2016, 12 (3), 1368–1379. 10.1021/acs.jctc.5b00992.

(105) Müller, C.; Schlauderer, G.; Reinstein, J.; Schulz, G. Adenylate Kinase Motions during Catalysis: An Energetic Counterweight Balancing Substrate Binding. Structure 1996, 4 (2), 147–156. 10.1016/S0969-2126(96)00018-4.

(106) Wu, Q.; Peng, Z.; Zhang, Y.; Yang, J. COACH-D: Improved Protein–Ligand Binding Sites Prediction with Refined Ligand-Binding Poses through Molecular Docking. Nucleic Acids Research 2018, 46 (W1), W438–W442. 10.1093/nar/gky439.

(107) Case, D. A.; Belfon, K.; Ben-Shalom, I.; Brozell, S.; Cerutti, D.; Cheatham III, T.; Cruzeiro, V.; Darden, T.; Duke, R.; Giambasu, G.; Gilson, M. K.; Gohlke, H.; Goetz, A.; Harris, R.; Izadi, S.; Izmailov, S.; Kasavajhala, K.; Kovalenko, A.; Krasny, R.; Kurtzman, T.; Lee, T.; LeGrand, S.; Li, P.; Lin, C.; Liu, J.; Luchko, T.; Luo, R.; Man, V.; Merz, K.; Miao, Y.; Mikhailovskii, O.; Monard, G.; Nguyen, H.; Onufriev, A.; Pan, F.; Pantano, S.; Qi, R.; Roe, D.; Roitberg, A.; Sagui, C.; Schott-Verdugo, S.; Shen, J.; Simmerling, C.; Skrynnikov, N.; Smith, J.; Swails, J.; Walker, R.; Wang, J.; Wilson, L.; Wolf, R.; Wu, X.; Xiong, Y.; Xue, Y.; York, D.; Kollman, P. AMBER20, 2020.

(108) Maier, J. A.; Martinez, C.; Kasavajhala, K.; Wickstrom, L.; Hauser, K. E.; Simmerling, C. ff14SB: Improving the Accuracy of Protein Side Chain and Backbone Parameters from ff99SB. J. Chem. Theory Comput. 2015, 11 (8), 3696–3713. 10.1021/acs.jctc.5b00255.

(109) Wang, L.-P.; Martinez, T. J.; Pande, V. S. Building Force Fields: An Automatic, Systematic, and Reproducible Approach. J. Phys. Chem. Lett. 2014, 5 (11), 1885–1891. 10.1021/jz500737m.

(110) Wang, L.-P.; McKiernan, K. A.; Gomes, J.; Beauchamp, K. A.; Head-Gordon, T.; Rice, J. E.; Swope, W. C.; Martínez, T. J.; Pande, V. S. Building a More Predictive Protein Force Field: A Systematic and Reproducible Route to AMBER-FB15. J. Phys. Chem. B 2017, 121 (16), 4023–4039. 10.1021/acs.jpcb.7b02320.

(111) Abraham, M. J.; Murtola, T.; Schulz, R.; Páll, S.; Smith, J. C.; Hess, B.; Lindahl, E. GROMACS: High Performance Molecular Simulations through Multi-Level Parallelism from Laptops to Supercomputers. SoftwareX 2015, 1–2, 19–25. 10.1016/j.softx.2015.06.001.

(112) Tribello, G. A.; Bonomi, M.; Branduardi, D.; Camilloni, C.; Bussi, G. PLUMED 2: New Feathers for an Old Bird. Computer Physics Communications 2014, 185 (2), 604–613. 10.1016/j.cpc.2013.09.018.

(113) Sousa da Silva, A.W.; Vranken, W. F. ACPYPE - AnteChamber PYthon Parser interfacE. BMC Res Notes 2012, 5 (1), 367. 10.1186/1756-0500-5-367.

(114) Ponzoni, L.; Polles, G.; Carnevale, V.; Micheletti, C. SPECTRUS: A Dimensionality Reduction Approach for Identifying Dynamical Domains in Protein Complexes from Limited Structural Datasets. Structure 2015, 23 (8), 1516–1525. 10.1016/j.str.2015.05.022.

(115) Honorato, R. V.; Trellet, M. E.; Jiménez-García, B.; Schaarschmidt, J. J.; Giulini, M.; Reys, V.; Koukos, P. I.; Rodrigues, J. P. G. L. M.; Karaca, E.; van Zundert, G. C. P.; Roel-Touris, J.; van Noort, C. W.; Jandová, Z.; Melquiond, A. S. J.; Bonvin, A. M. J. J. The HADDOCK2.4 Web Server: A Leap Forward in Integrative Modelling of Biomolecular Complexes. Nature Protocols 2024, in press.

(116) Wang, J.; Wolf, R. M.; Caldwell, J. W.; Kollman, P. A.; Case, D. A. Development and Testing of a General Amber Force Field. Journal of computational chemistry 2004, 25 (9), 1157–1174.

