## Supplemental Material for "Predicting binding events in very flexible, allosteric, multi-domain proteins"

### Supplementary Figures

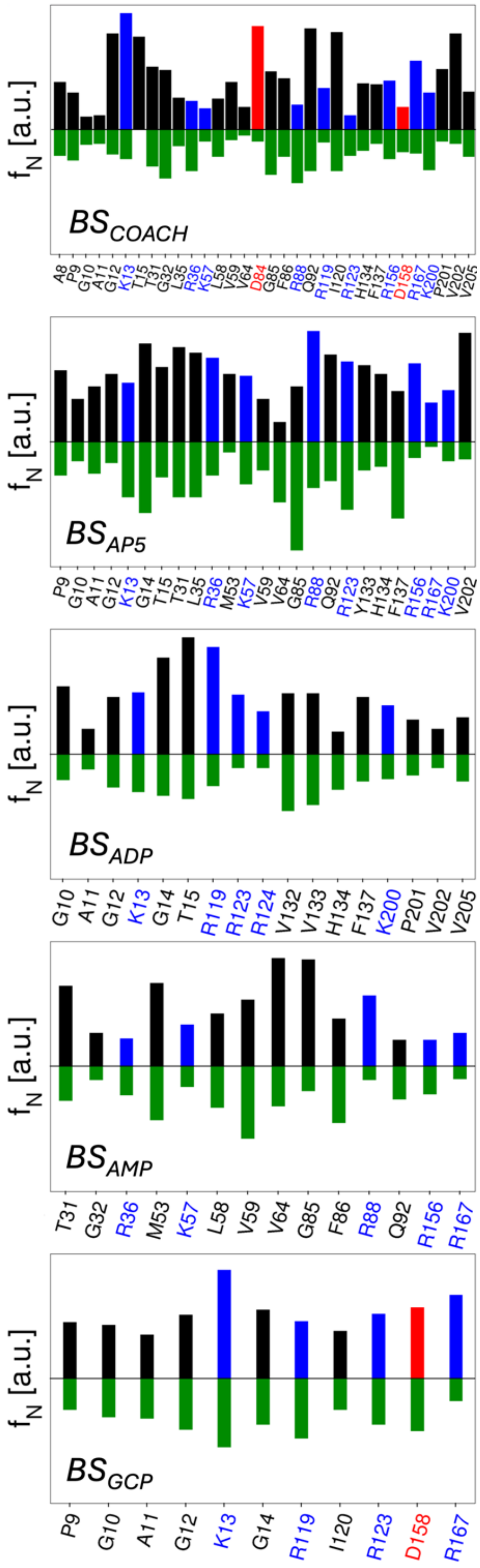


**Figure S1**. Frequency of holo-like conformations sampled by each residue lining the BS described in the main text. Bars pointing upwards refer to gEDES and are colored according to the amino acid type (red: negatively charged; blue: positively charged; black: neutral), while green bars pointing downwards refer to the unbiased simulation. A holo-like conformation is counted when the RMSD of the residue (calculated on all non-hydrogenous atoms after alignment of the whole BS) is lower than the arbitrary threshold defined for each amino acid in **Table S6** (and always lower than 2.5 Å).

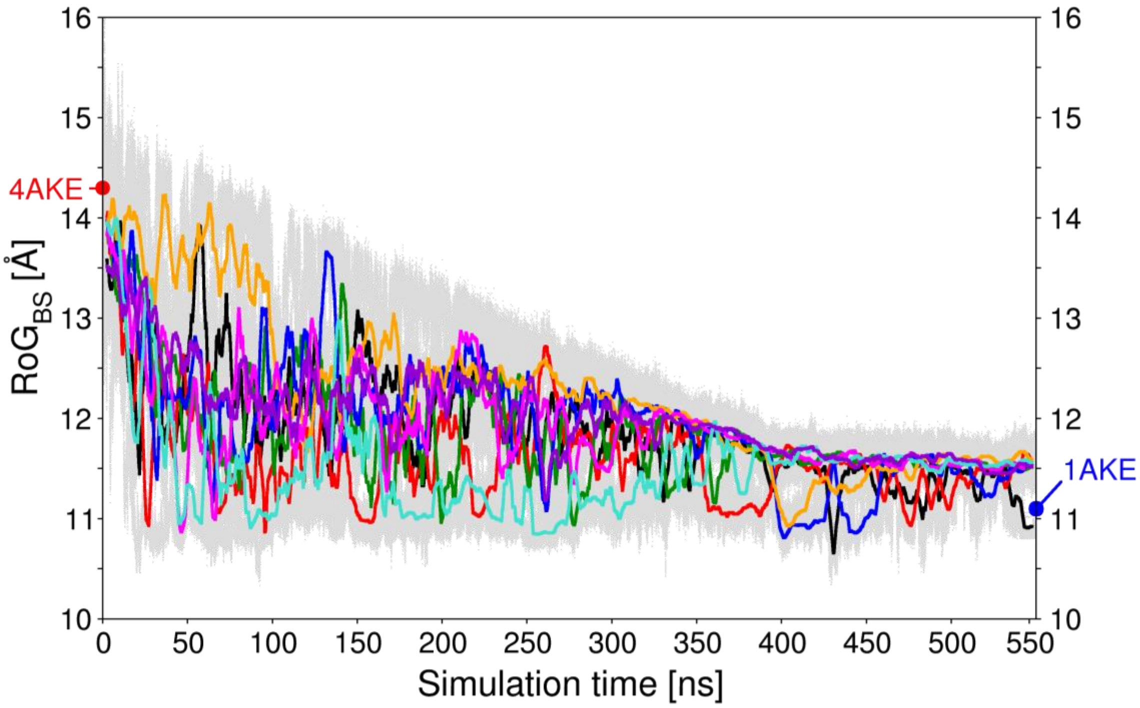


**Figure S2.** Plots of the radius of gyration (RoG) calculated for $BS_{COACH}$ (backbone atoms) along the 8 replicas of the gEDES trajectory discussed in the main text. Values of the cumulative trajectory from all replicas are represented as grey points, while the running averages (of length 2500 points, corresponding to 50 ns) calculated on each replica are represented by colored lines. $RoG_{BS}$ values for the experimental apo (14.3 Å; PDB ID: 4AKE) and holo (11.1 Å; PDB ID: 1AKE) structures are indicate by a red and a blue sphere, respectively. The standard deviations of $RoG_{BS}$ for each replica amounted to 3, 1.8, 0.8, 3, 1.8, 0.9, 0.9, 0.8 % of the corresponding average.

### Supplementary Tables

| Binding region | ResIDs |
|---|---|
| **BS1$_{COACH}$** | **A8** P9 G10 A11 G12 K13 T15 R119 K200 P201 V202 V205 |
| **BS2$_{COACH}$** | P9 G10 A11 G12 K13 T15 T31 G32 L35 R36 K57 L58 V59 V64 G85 F86 R88 Q92 R119 I120 R123 H134 F137 R156 R167 K200 V202 |
| **BS3$_{COACH}$** | P9 T31 G32 L35 R36 K57 L58 V59 V64 **D84** G85 F86 R88 Q92 R123 **D158** R167 |
| **BS$_{COACH}$** | **A8** P9 G10 A11 G12 K13 T15 T31 G32 L35 R36 K57 L58 V59 V64 **D84** G85 F86 R88 Q92 R119 I120 R123 H134 F137 R156 **D158** R167 K200 P201 V202 V205 |
| **NC** | T31 G32 L35 R36 K57 L58 V59 V64 D84 G85 F86 R88 Q92 |
| **LC** | A8 P9 G10 A11 G12 K13 T15 R119 I120 R123 H134 F137 R156 D158 R167 |
| **BS$_{AP5}$** | P9 G10 A11 G12 K13 **G14** T15 T31 L35 R36 **M53** K57 V59 V64 G85 R88 Q92 R123 **Y133** H134 F137 R156 R167 K200 V202 |
| **BS$_{ADP}$** | G10 A11 G12 K13 **G14** T15 R119 R123 **R124** **V132** **Y133** H134 F137 K200 P201 V202 V205 |
| **BS$_{AMP}$** | T31 G32 R36 **M53** K57 L58 V59 V64 G85 F86 R88 Q92 R156 R167 |
| **BS$_{GCP}$** | P9 G10 A11 G12 K13 **G14** R119 I120 R123 D158 R167 |

**Table S1.** Residues lining the different binding regions considered in this work. Bolded labels in BS$_{COACH}$ indicate residues not retrieved in any of the experimental binding sites. Bolded labels in BS$_{AP5, ADP, AMP, GCP}$ indicate residues not included in the definition of BS$_{COACH}$. Positively (R, K) and negatively (D) charged residues are colored blue and red, respectively.

| Domain | $C_\alpha$–RMSD |
|---|---|
| NMP | 1.6 |
| LID | 0.6 |
| CORE | 1.6 |

**Table S2**. Inter-domain RMSD between experimental apo (PDB ID: 4AKE) and holo (PDB ID: 1AKE) structures. RMSD refers to $C_\alpha$ atoms after domain alignment.

| CV | ResIDs | CV width |
|---|---|---|
| **RoG$_{BS}$** | 8 9 10 11 12 13 15 31 32 35 36 57 58 59 64 84 85 86 88 92 119 120 123 134 137 156 158 167 200 201 202 205 | 0.3 |
| **CIP$_1$** | 31 32 35 36 57 58 59 64 84 85 86 88 92 167<br>8 9 10 11 12 13 15 119 120 123 134 137 156 158 200 201 202 205 | 7.3 |
| **CIP$_2$** | 9 10 32 35 36 57 58 59 119 120 123 134 137 156 158 167<br>8 11 12 13 15 31 64 84 85 86 88 92 200 201 202 205 | 4.0 |
| **CIP$_3$** | 8 9 10 11 57 58 59 85 86 88 92 119 120 158 167 200 205<br>12 13 15 31 32 35 36 64 84 123 134 137 156 158 201 202 | 8.6 |
| **cRD$_{NCc}$** | 84 88<br>36 57 | 0.9 |
| **cRD$_{NCo}$** | 31 32 35 58 59 64<br>85 86 92 | 4.1 |
| **cRD$_{LCc}$** | 119 123 134 156<br>13 167 | 2.5 |
| **cRD$_{LCo}$** | 8 9 10 11 12 15<br>120 137 158 | 6.2 |

**Table S3**. Implementation details of CVs used in gEDES approach. Columns 1-3 refer respectively to CV's name, IDs of the selected residues, and to the width of energy hills. Residue groups defining CIP and cRD variables are reported on different rows.

| BS | $RoG_{X\text{-}ray}^{apo}$ [Å] | $RoG_{X\text{-}ray}^{holo}$ [Å] | ΔRoG (%) |
|---|---|---|---|
| $BS_{AP5}$ (NC+LC) | 14.3 | 11.1 | 22 |
| $BS_{ADP}$ (LC) | 10.4 | 7.7 | 26 |
| $BS_{AMP}$ (NC) | 11.5 | 8.7 | 24 |
| $BS_{GCP}$ (LC) | 8.3 | 7.8 | 6 |

**Table S4**. Percentage of RoG variation, calculated on backbone atoms, for the different BS investigated here, between the apo protein (PDB ID: 4AKE) and the relative complex. Complex structures used for defining $BS_{AP5}$, $BS_{GCP}$, $BS_{ADP}$ and $BS_{AMP}$ are identified respectively by PDB IDs: 1AKE, 6F7U, 2ECK, 1ANK. For each BS selection, its position referred to NC/LC interfaces is also reported.

| | | 1AKE | 2ECK | 1ANK |
|---|---|---|---|---|
| **Protein ($C_\alpha$)** | 1AKE | - | | |
| | 2ECK | 0.3 | - | |
| | 1ANK | 0.5 | 0.5 | - |
| | 6F7U | 6.3 | 6.4 | 6.3 |
| **$BS_{AP5}$ (noh)** | 1AKE | - | | |
| | 2ECK | 0.8 | - | |
| | 1ANK | 0.4 | 0.8 | - |
| | 6F7U | 5.4 | 5.4 | 5.4 |
| **$BS_{ADP}$ (noh)** | 1AKE | - | | |
| | 2ECK | 0.4 | - | |
| | 1ANK | 0.8 | 0.7 | - |
| | 6F7U | 3.4 | 3.4 | 3.3 |
| **$BS_{AMP}$ (noh)** | 1AKE | - | | |
| | 2ECK | 0.4 | - | |
| | 1ANK | 0.8 | 0.8 | - |
| | 6F7U | 5.3 | 5.2 | 5.3 |
| **$BS_{GCP}$ (noh)** | 1AKE | - | | |
| | 2ECK | 2.2 | - | |
| | 1ANK | 2.3 | 0.8 | - |
| | 6F7U | 2.1 | 2.2 | 2.2 |

**Table S5.** RMSD matrix (Å) among different experimental holo complexes for the whole protein and for the putative regions, after alignment of the selected region. Binding regions have been defined from protein structure in the experimental complex and enclose residues within 3.5 Å from the ligand. Complex structures used for defining $BS_{AP5}$, $BS_{ADP}$, $BS_{AMP}$ and $BS_{GCP}$, are identified respectively by the structures with PDB IDs: 1AKE, 2ECK, 1ANK and 6F7U.

| Residue | $RMSD_{cutoff}$ (Å) |
|---|---|
| A | 1.9 |
| R | 2.5 |
| N | 2.5 |
| D | 2.5 |
| C | 2.2 |
| Q | 2.5 |
| E | 2.5 |
| G | 1.7 |
| H | 2.5 |
| I | 2.5 |
| L | 2.5 |
| K | 2.5 |
| M | 2.5 |
| F | 2.5 |
| P | 2.3 |
| S | 2.5 |
| T | 2.5 |
| W | 2.5 |
| Y | 2.5 |
| V | 2.4 |

**Table S6.** Adaptive cut-off RMSD used to identify bound-like residue conformations. Values are calculated considering the number of side-chain non-hydrogenous atoms ($N_{heavy}$) and torsional angles ($N_{torsion}$) of each residue, according to the formula $RMSD_{cutoff}$ (Å) = 1.5 + 0.2 ($N_{heavy}$) + 0.3 ($N_{torsion}$) and a maximum cut-off set to 2.5 Å.
